# Model discovery approach enables non-invasive measurement of intra-tumoral fluid transport in dynamic MRI

**DOI:** 10.1101/2023.08.28.554919

**Authors:** Ryan T. Woodall, Cora C. Esparza, Margarita Gutova, Maosen Wang, Jessica J. Cunningham, Alexander B. Brummer, Caleb A. Stine, Christine C. Brown, Jennifer M. Munson, Russell C. Rockne

**Affiliations:** Division of Mathematical Oncology and Computational Systems Biology, Department of Computational and Quantitative Medicine, Beckman Research Institute, City of Hope National Medical Center; 1500 E Duarte Rd, Duarte, CA 91010, USA; Fralin Biomedical Research Institute, Virginia Institute of Technology at Virginia Tech Carilion, Virginia Tech, 4 Riverside Circle, Roanoke, VA 24016, USA; Department of Stem Cell Biology and Regenerative Medicine Beckman Research Institute, City of Hope National Medical Center; 1500 E Duarte Rd, Duarte, CA 91010, USA; Department of Physics and Astronomy, College of Charleston; 66 George Street, Charleston, SC 29424, USA; Hematology & Hematopoietic Cell Transplantation, Beckman Research Institute, City of Hope National Medical Center; 1500 E Duarte Rd, Duarte, CA 91010, USA; Department of Immuno-Oncology, Beckman Research Institute, City of Hope National Medical Center; 1500 E Duarte Rd, Duarte, CA 91010, USA

**Author notes:** Corresponding authors: Ryan T. Woodall, Jennifer M. Munson, Russell C. Rockne.

## Abstract

Dynamic contrast-enhanced magnetic resonance imaging (DCE-MRI) is a routine method to non-invasively quantify perfusion dynamics in tissues. The standard practice for analyzing DCE-MRI data is to fit an ordinary differential equation to each voxel. Recent advances in data science provide an opportunity to move beyond existing methods to obtain more accurate measurements of fluid properties. Here, we developed a localized convolutional function regression that enables simultaneous measurement of interstitial fluid velocity, diffusion, and perfusion in 3D. We validated the method computationally and experimentally, demonstrating accurate measurement of fluid dynamics *in situ* and *in vivo*. Applying the method to human MRIs, we observed tissue-specific differences in fluid dynamics, with an increased fluid velocity in breast cancer as compared to brain cancer. Overall, our method represents an improved strategy for studying interstitial flows and interstitial transport in tumors and patients. We expect that it will contribute to the better understanding of cancer progression and therapeutic response.

**One-Sentence Summary:** A physics-informed computational method enables accurate and efficient measurement of fluid dynamics in individual patient tumors and demonstrates differences between tissues.

## Main Text

Interstitial fluid transport is intrinsically linked to the movement of drugs, nutrients, and cells in tissues and is therefore especially important for understanding cancer physiology. Cancers in any tissue develop aberrant transport that can affect the movement of drugs into and through tumors. Interstitial fluid flow, in particular, has been identified as a driving force in tumor cell invasion into surrounding healthy tissue (*1–4*). Interstitial flow can also change the surrounding microenvironment, activating fibroblasts (*5, 6*), directing immune cell behavior (*7, 8*), and inducing angiogenesis (*9, 10*). As such, understanding interstitial flows and interstitial transport is vital to understanding disease progression and therapeutic application strategies (*11, 12*), though measuring the interstitial fluid flow field non-invasively has remained a challenge, particularly in situ and in 3D.

Contrast-enhanced magnetic resonance imaging (MRI) has provided clinical benefit for cancer treatment for many decades (*13*). Given its non-invasive nature and ability to image soft tissues, which are usually difficult to see with other imaging technologies, MRI has become a staple of diagnosing, planning, and providing prognosis in several cancer settings, especially in brain (*14–16*). Importantly, MRI allows acquisition of physiologically relevant information in both space and time, and therefore represents one of the best methods for studying dynamics *in situ* and in 3D. Over two decades ago, a version of MRI called dynamic contrast-enhanced (DCE-MRI) was developed to allow the study of tissue vasculature (*17*). An DCE-MRI experiment measures signal enhancement due to the presence of an intravenously injected paramagnetic contrast agent in the region of interest. In practice, a series of images are acquired over time, capturing the signals before, during and after the contrast agent reaches the target tissue, allowing for both visualization and quantification of vasculature in space and in time. Thus, DCE-MRI may offer an improved strategy for studying interstitial flows and interstitial transport.

Existing data processing approaches for physical interpretation of contrast enhancement data from DCE-MRI use ordinary differential equation (ODE) models applied to individual voxels, though this method largely fails to incorporate the rich spatial data provided by the modality (*18*). Few methods exist for quantifying the spatially varying effective diffusion coefficient of contrast agent in DCE-MRI (*19, 20*), and similarly few attempt to quantify and investigate interstitial fluid flow (i.e. advection) within the tumor (*21–23*). While there do exist MRI sequences which directly measure interstitial fluid velocity, they struggle with separating vascular flow from interstitial flow, and require additional imaging sequences to be applied in the clinic, significantly increasing the time a patient is in the scanner (24, 25). Given that DCE-MRI is become a standard clinical practice, there is growing opportunity to utilize it for studying interstitial flow, without the need for specialized sequences (26, 27). Here, we develop a strategy that overcomes these limitations by integrating DCE-MRI with a method developed for the discovery of equations governing time-series data through the concept of function-space regression, known in data sciences as sparse identification of nonlinear dynamics (SINDy) (*28*). Given that the time- and space-resolved data obtained by DCE-MRI is often noisy (*29*), we modified weak SINDy, a method that leverages the weak form of governing equations to provide an efficient and accurate method for simultaneous model discovery and parameter estimation from noisy data (*30*). This method bypasses the use of discrete approximations of derivatives on noisy or sparse data, which are used by SINDy (*28*), or iterative gradient descent methods used in modern DCE-MRI model inversion which require estimating derivatives of the objective function with respect to the data (*21–23*). The key insight from weak SINDy is to integrate raw data with derivatives of a known basis function, selected by the user to be optimal for the problem at hand (31). This method serves to smooth the data and results, instead of taking a finite-difference approximation of noisy data. We validated the method on simulated data, as well as experimentally in a hydrogel model. Further, we demonstrated that our method works well with data obtained *in vivo* using a mouse xenograft model, and showed the clinical potential by examining fluid dynamics in human glioblastoma brain tumors and breast cancer data sets (*32, 33*). Taken together, our approach to DCE-MRI data processing offers a fast and noise-robust strategy for studying interstitial flows and general fluid transport in space and time.

## Results

### Tailoring weak SINDy for processing DCE-MRI data

Here, we started from weak SINDy and customized it to handle the partial differential equations (PDEs) relevant to DCE-MRI, and perform localized convolutional function regression (LCFR) to recover spatially localized fluid transport parameters (Fig. 1, A). Briefly, we assumed that the dynamics of contrast agent transport are governed by an advection-diffusion-reaction PDE, with vascular input forcing function (VIF), and an unmixed enhancing plasma-compartment (*vp*), consistent with extended Tofts-Kety intra-voxel transport (*K* ^*trans*^,*k*^*ep*^),advective 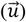 and diffusive (*D*) inter-voxel transport:

**Figure 1:**
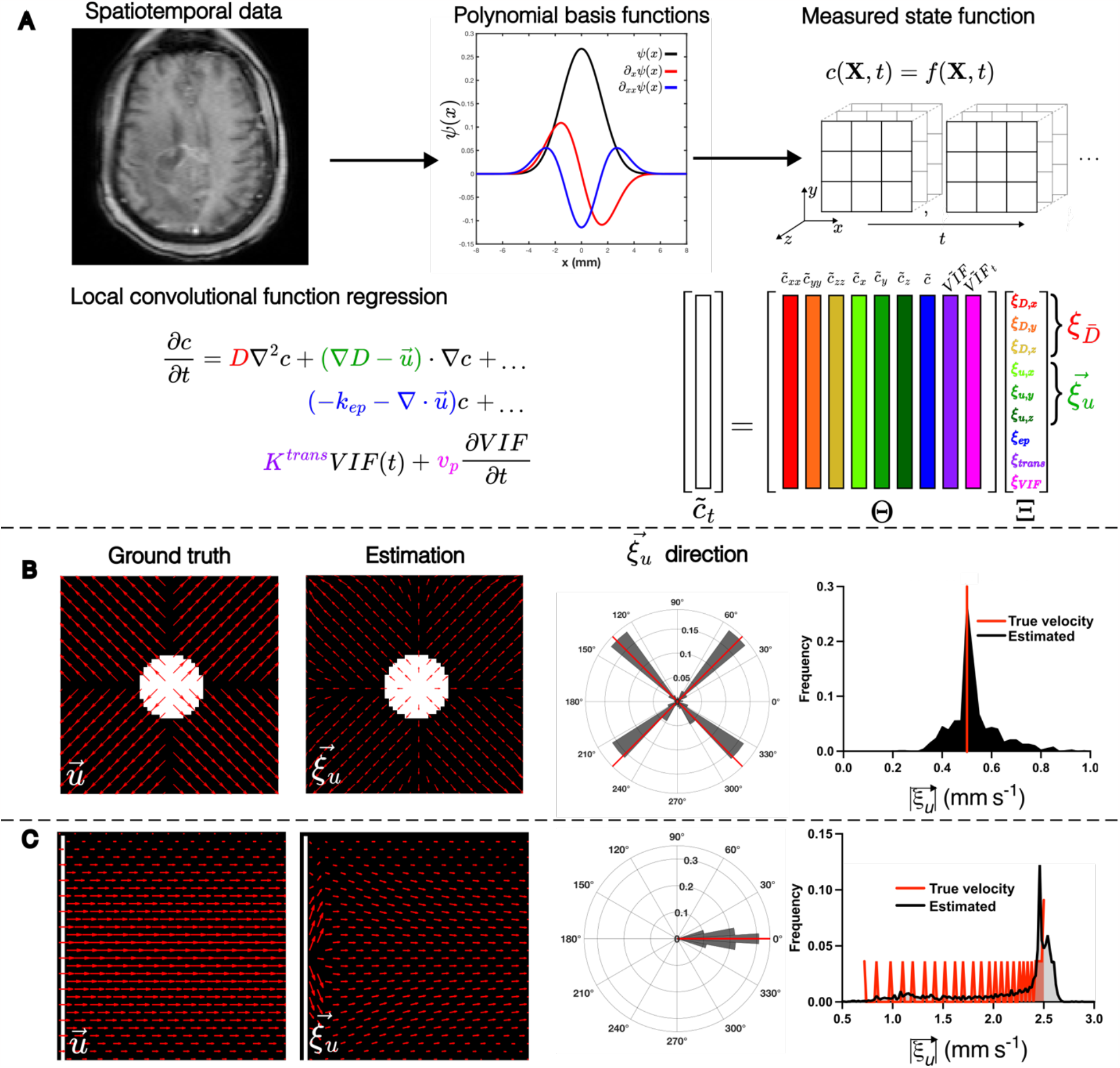
LCFR Methodology and validation with in silico phantoms. (A) Methodology of Localized Convolutional Function Regression (LCFR), wherein spatiotemporal contrast agent concentration data is convolved with a smooth basis-function and its derivatives, divided into 3x3x3 windows. The coefficients of the factored transport PDE are then solved for using linear regression. (B) Validation of LCFR coefficient 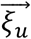 on a divergent flow field (initial condition, white) with spatially invariant diffusion, and the true direction and magnitude of velocity denoted in red bars. (C) Validation of CLFR on a Poiseuille shear flow field with spatially invariant diffusion (initial condition, white), and the true direction and magnitude of velocity denoted in red bars.

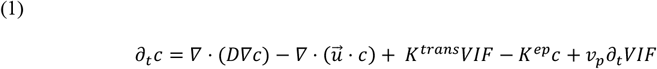

We then projected the concentration of contrast agent, c(***X***, *t*), onto smooth, 4D polynomial basis functions Ψ (***X***, *t*), to perform a spatially localized model regression of the form:

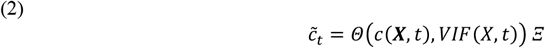

where 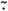 indicates convolution of the data onto basis functions or their derivatives, denoted as 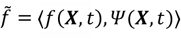 and 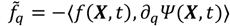, respectively (Supplemental Equation 21-22). The model regression problem was then formulated on a (x, y, z) 3x3x3 window where a weak derivative of the data is performed via the basis functions. The matrix Θ(*c*(***X***, *t*), *VIF* (***X***, *t*)) is populated with polynomial projections of the data and corresponding weak derivatives,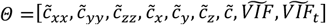. The model regression problem was then solved locally in space, for all time, to recover the vector Ξ, which consists of coefficients of the governing equation (1) as, Ξ = [ξ_*D,x*_, ξ _*D,y*_, ξ _*D,z*_, ξ _*u,x*_, ξ _*u,y*_, ξ _*u,z*_, ξ _*ep*_, ξ _*trans*_, ξ _*VIF*_]^*T*^. A mapping between the coefficients and the terms of the full expansion of equation (1) can be found in the Supplement (S14, S15). This methodology allows for the most likely transport model coefficients to be obtained locally within the original image context, and does so efficiently with simply regression. Below, we assess the accuracy of this method using multiple methods.

### LCFR accurately estimates transport parameters from simulated data

To validate our approach, we first examined whether our methodology is able to recover local PDE coefficients *in silico* from forward model simulations. An advection-diffusion model was used to generate two flow patterns (Supplemental Table 1). The first model we considered is a radially diverging flow field (Fig. 1, B), which was chosen because it simulates the outward flow typically expected of a high-pressure tumor, and will test the method’s ability to distinguish advective and diffusive dispersion. The second model we implemented was a Poiseuille shear flow (Fig. 1, C), a standard case of incompressible flow in a pipe or blood vessel, with a well-studied spatially varying velocity field. In both cases, LCFR accurately recovers the direction and magnitude of the flow field after addition of 0.1% noise (% maximum signal, for stabilization of the method), with root mean squared error (RMSE) of 1.02E-1 mm/s for divergent flow, and RMSE = 4.63E-1 mm/s for Poiseuille flow. Of note, we removed edge effects due to the Gibbs phenomenon from the visualization and analysis. To test the ability of the method to identify the transport rate constant *K*^*trans*^ and vascular volume fraction *v*_*p*_, we simulated the extended Tofts-Kety dynamics with spatially varying perfusion and 0.1% noise, *K*^*trans*^ and constant vascular volume fraction, *v*_*p*_ = 5.00E-2. In this scenario, *K*^*trans*^ was accurately estimated by the coefficient *ξ*_*trans*_ (RMSE = 1.54E-1 1/s), while the measurement of the plasma volume fraction *v*_*p*_ (RMSE = 3.89E-1) was observed to be influenced by the transport rate Ktrans (**Table S1, Figure S3**). Taken together, our *in silico* validation demonstrates the ability of our method to accurately capture both the direction and magnitude of different fluid velocity fields. We showed that sharp changes in the field can be smoothed out by the basis functions, though the resulting fields are consistent with the true velocity fields. We also demonstrate measurement of Tofts-Kety-like dynamics, wherin *ξ*_*trans*_ accurately estimates *K*^*trans*^. This gave us confidence to further validate LCFR *in vitro, in vivo* and using clinical data.

### LCFR accurately measures the mean contrast velocity in porous hydrogel

We next validated the methodology *in vitro*, by administering a bolus of contrast agent on top of porous hydrogel with a pressure head forcing the contrast agent through the gel (Fig. 2, A). DCE-MRI was acquired at 30 second time intervals, and signal intensity was converted to contrast agent concentration using T1-mapping. The LCFR was applied to the resulting concentration field. We manually estimated the velocity of the contrast agent front (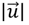 =1.14E-3 mm/s) (Fig 2. B) and compared it to the LCFR-estimated 3D velocity 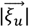 within the hydrogel (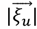 =1.32E-3 ± 8.20E-4 mm/s) (Fig 2, B-E). The mean measured diffusivity within the gel 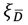was measured to be 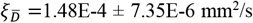 (N = 3). This analysis was performed on multiple replicates, and the mean error associated with the method was 15.2% ± 1.08% (N = 3) (Fig 2. E). Overall, these results illustrate that using LCFR results in accurate estimation of 3D velocity and diffusivity in a controlled system, where there are reliable methods for estimating the true velocity rate and literature-characterized diffusivity. From these findings, we began investigating the results of our methodology *in vivo*.

**Figure 2:**
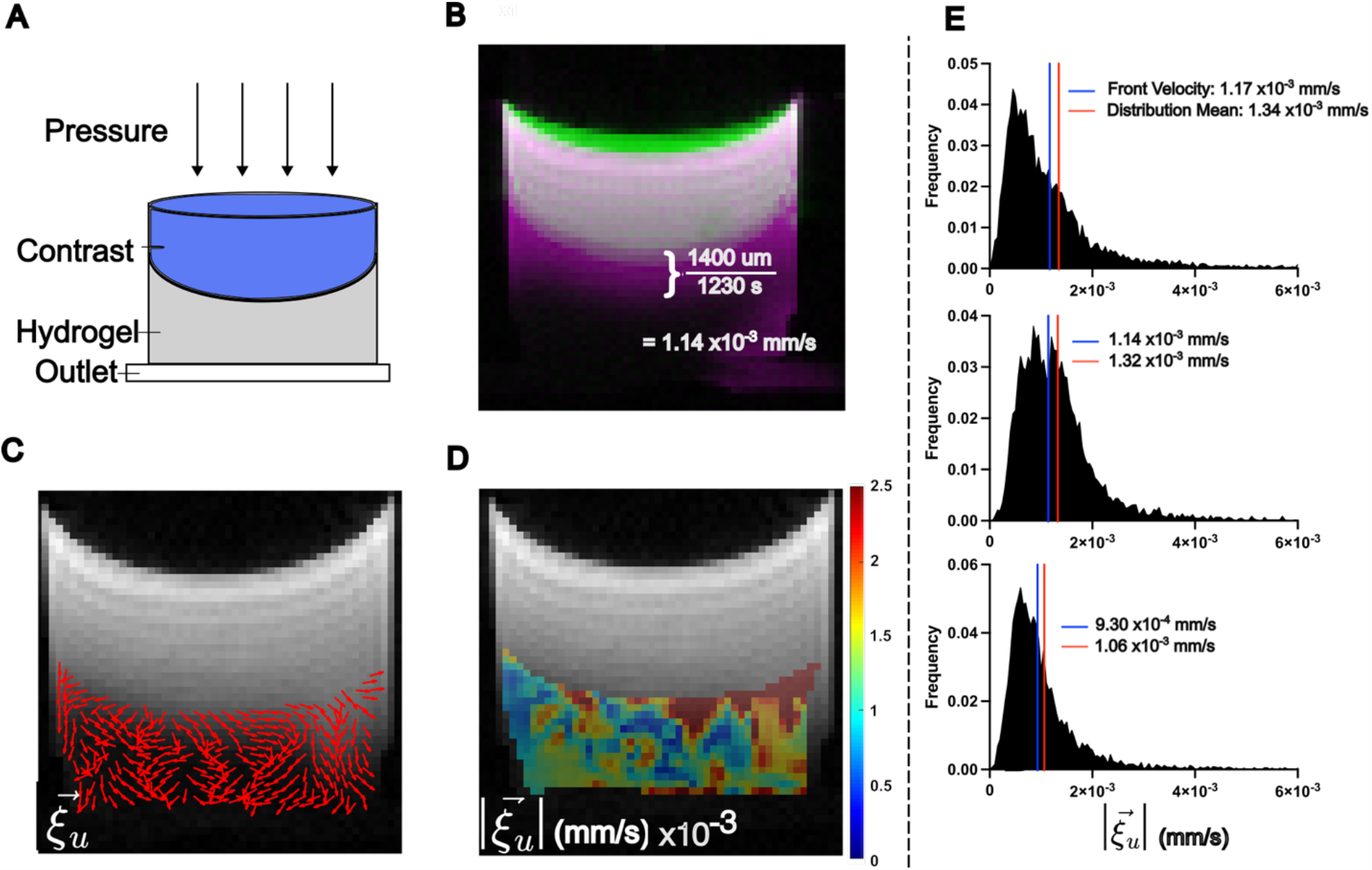
LCFR predicts interstitial fluid velocity in hydrogel phantoms. (A) Experimental setup, wherein a bolus of contrast agent is administered onto of a porous hydrogel and drains through due to a hydraulic pressure head. (B) Method of estimating the mean flow velocity of the contrast agent front, using the difference between the initial contrast location (green), and final contrast location (pink), resulting in an estimated contrast agent velocity of 1.14E-3 mm/s. (C) The estimated in-plane interstitial flow velocity direction overlaid on the final T1-weighted image. (D) The local 3D magnitude of interstitial flow velocity within the hydrogel. (E) Histograms of three replicate trials, comparing the 3D magnitude of flow velocity within the hydrogel, with blue line indicating the estimated contrast front velocity, and red line indicating the mean velocity of the distribution as measured by LCFR.

### LCFR-measured perfusion predicts Evans Blue leakage into tumors in vivo

Encouraged by the results from *in silico* and *in vitro* analysis, we subsequently validated the method *in vivo*. Six mice (N = 6) implanted with a mouse glioma cell line (GFP-GL261) into the brain underwent DCE-MRI with isotropic spatial resolution of 0.2 mm, at 7 and 14 days post-implantation. After imaging on day 14, the mice were injected with Evans Blue to quantify perfusion, and brains were harvested and stained for visual comparison to LCFR outputs (Fig. 3, A through D). For comparison to histology, the estimated perfusion *ξ*_*trans*_ is overlaid on the central tumor slice of the T1-weighted image (Fig. 3E). The mean value of ξ_*trans*_ was then compared to the mean area coverage of Evans Blue after venous infusion, and is observed to be positively correlated with the mean tumor perfusion rate constant, *ξ*_*trans*_ across all mice (r = 0.507, P = 0.038, N = 17, two-tailed Pearson correlation) (Fig. 3, F). The resulting 3D interstitial velocity field, 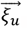 is displayed over the native post-contrast T1 weighted volume, highlighting that LCFR is readily applied on 4D data (Fig 3, G). The measured velocity 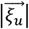 remained constant from 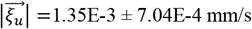 on day 7, to 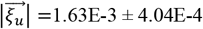 mm/s on day 14 (P = 0.218, N = 6, two-tailed Wilcoxon test) (Fig 3, H). The fluid rate transfer constant, *ξ*_*trans*_increased from 2.70E-2 ± 8.65E-3 1/s on day 7, to 5.08E-2 ± 1.84E-2 1/s on day 14 (P = 0.063, N = 6, two-tailed Wilcoxon test) (Fig 3, I). These results demonstrate that LCFR can non-invasively detect individual variation of perfusion and flow, as validated with histology.

**Figure 3:**
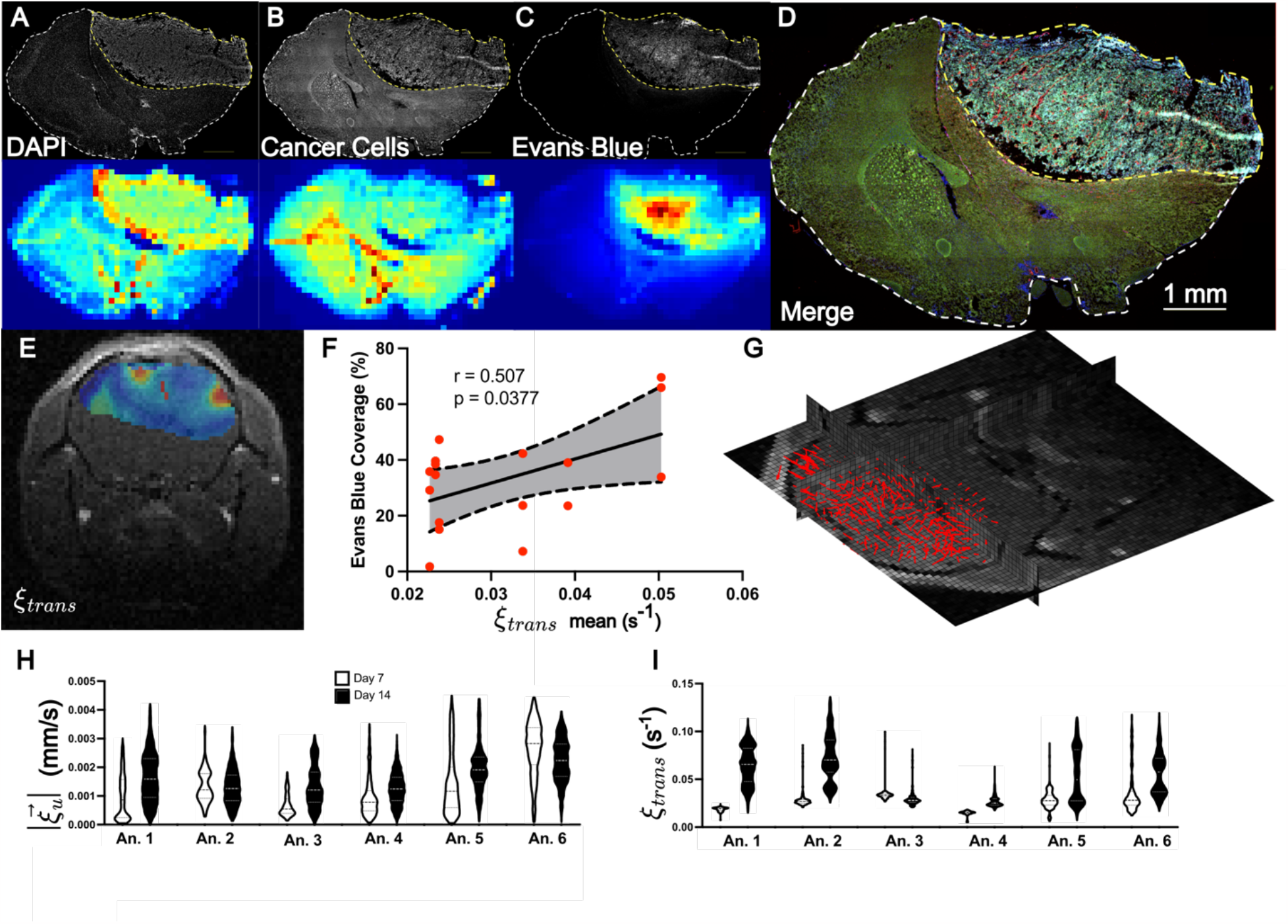
LCFR-measured perfusion is correlated to Evans Blue coverage in vivo. (A through D) Representative coronal IHC stains through the central tumor slice (top row), and MR resolution-matching intensity projection (bottom row), consisting of DAPI (A), GFP-expressing GL261 cells (B), Evans Blue (C). (D) Merge of all IHC demonstrating tumor heterogeneity. (E) Estimated perfusion field, *ξ*_*trans*_ overlaid on post-contrast T1-weighted image. (F) Scatter plot depicting the correlation and 95% confidence interval of linear regression between mean tumor perfusion as measured by DCE-MRI (*ξ*_*trans*_ x-axis), and Evans Blue Coverage (mean tumor stain intensity, y-axis), for n = 17 histology slices. (G) 3D velocity vector field of estimated interstitial velocity 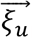 overlaid on the 3D T1 post-contrast volume. (H and I) Violin plots depicting the estimated velocity magnitude (H) and perfusion (I) for 6 animals imaged 7 and 14 days after tumor implantation.

**Figure 4:**
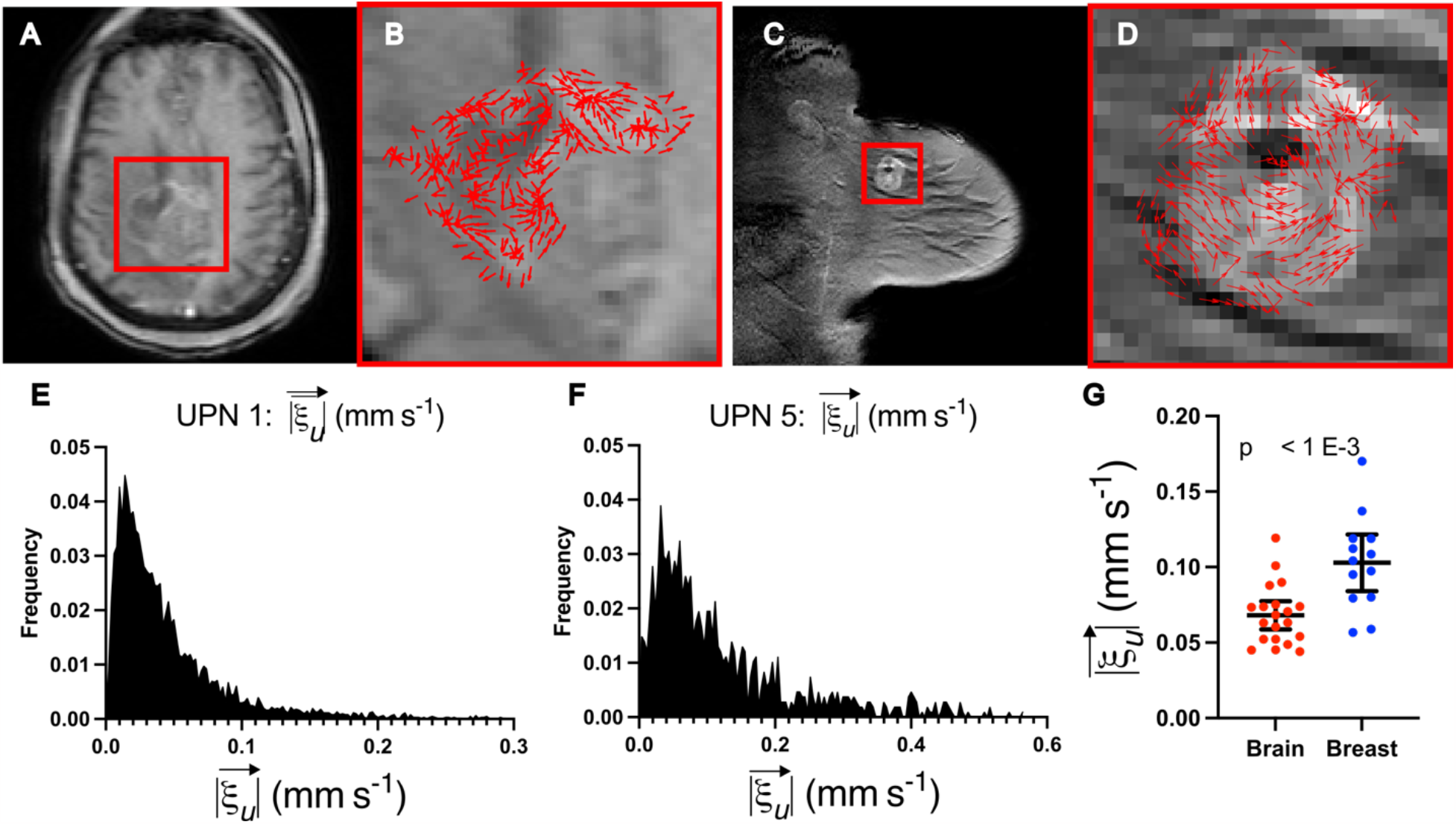
LCFR captures differences in fluid transport between breast and brain cancers. (A and B) Representative post-contrast T_1_-weighted image of central slice of residual glioblastoma two weeks after resection surgery. (B) Detail of tumor and resection cavity, with overlay of the estimated velocity direction, 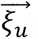 (C and D) Representative post-contrast T_1_-weighted image of untreated primary breast cancer lesion. (D) Detail of the enhancing tumor, with overlay of the with overlay of the estimated velocity direction, 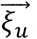 (E) Distribution of the in-plane velocity of the entire enhancing glioblastoma and resection cavity (mean = 5.23E-1 ± 5.10E-1). (F) Distribution of the in-plane velocity of the entire enhancing breast tumor (mean = 1.19E-1 ± 1.06E-1). (G) Mean fluid velocities for brain (mean = 6.81E-2 ± 1.99E-2 mm/s, N = 20) and breast data (mean = 1.03E-1 ± 3.10E-2, N = 13), with whiskers indicating mean and 95% confidence interval.

### LCFR-measured interstitial fluid velocity varies between breast and brain cancers

To investigate the differences in interstitial fluid flow between cancers in different tissues, we applied LCFR to clinical DCE-MRI in a cohort of post-resection treatment naive glioblastoma patients, and a cohort of treatment-naive breast cancer patients. In the glioblastoma cohort, 20 patients with recurrent disease who underwent DCE-MRI imaging between January 2020 and July 2022 were selected from the radiology records at City of Hope National Medical Center. The glioblastoma patients underwent DCI-MRI on average 32.3 days after resection and prior to receiving additional therapy. The breast cancer cohort consisted of 13 treatment-naïve patients from the Quantitative Imaging Network BREAST-02 study (*33*). In each dataset, the enhancing lesion was manually segmented, and LCFR was run to investigate the population fluid dynamical profile. In these cohorts, we measured the flow velocity, 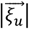, in breast tumors (1.03E-1 ± 3.10E-2 mm/s) to be significantly faster than that in brain tumors (6.81E-2 ± 1.99E-2 mm/s) (P < 0.001, N_brain_ = 20, N_breast_ = 13, two-tailed Mann Whitney test). These preliminary results indicate that cancers of different organs and cellular origins may present with different flow profiles, and may provide novel methods for explaining differences in disease progression and treatment response. These results warrant further exploration, which is outside the scope of this initial reporting of our novel methodology.

## Discussion

Interstitial fluid transport plays a key role in many processes connected with cancer physiology, tumor microenvironment, tumor immune response, as well as the response of cancer to treatment. However, physical and physiological properties of interstitial flows and transport remain difficult to study and understand. Here, we describe a novel methodology for analyzing DCE-MRI data, which allows for accurate, non-invasive measurement of fluid dynamics in living tissues. The methodology leverages the rich spatial and temporal data provided by DCE-MRI to enhance our understanding of tissue fluid dynamics. We have validated our approach using synthetic and experimental data, both *in vitro* by following the flow of contrast agent in a hydrogel, and *in vivo* by using a mouse model of glioblastoma. We also demonstrate application of LCFR to human MRI data routinely collected in the clinic from patients with either breast or brain cancer.

LCFR was able to accurately measure the mean 3D flow velocity of contrast agent forced through a hydrogel, and yielded mean diffusivity in the gel, 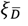, consistent with diffusivity measurements reported previously (*34, 35*). However, if we compare our results obtained using LCFR to process DCE-MRI mouse data (1.35E-3 ± 7.04E-4 mm/s) with results from recent studies that used phase-contrast imaging to directly measure fluid velocity within tumors (1.10E-1 ± 5.5E-4 mm/s and 1.10E-1 ± 5.5E-4 mm/s) (*24, 25*), we find some discrepancy, likely due to contribution from vascular flow, which could not be disambiguated from tissue interstitial fluid flow in the phase-contrast methods resulting in higher values (*24, 25,36,37*). In the present study, the bolus arrival time is corrected for in each voxel, thus minimizing the contribution of vascular velocity in the total velocity field. Additionally, we demonstrate (Supplemental Equation 14) that the spatial gradient of diffusion is structurally unidentifiable from true advective transport and may thus artificially increase the apparent velocities observed in past studies, and others directly measuring advective transport. These confounding factors may contribute to higher apparent velocities measured by methods which utilize higher temporal resolution data, or do not account for diffusive and advective transport separately, thus explaining the observed discrepancy.

Our methodology has several advantages over traditional model inversion methods used to parameterize DCE-MRI data, foremost that it is readily applied in native 4D, instead of individual z-slices, allowing for simultaneous estimation of both inter- and intra-voxel fluid transport parameters. Moreover, LCFR is robust to noise due to the use of smooth polynomial basis functions. Our method is highly efficient, utilizing the fast Fourier Transform for convolution with basis functions, and linear regression for the recovery of local PDE coefficients. Finally, our method allows for a direct calculation of parameters of a PDE from the original data, agnostic of spatial and temporal resolutions, and without the need for iterative forward-PDE solutions required for PDE inverse problems (*38, 39*).

While these advantages are useful for this type of data, they do come with trade-offs which limit the performance of the method. For example, our method uses a pre-defined library of functions to characterize the dynamics (Θ), instead of an extensive library of hypothetical functions and polynomial combinations of partial derivatives. This is largely because neither the higher-order and spatial cross-derivatives, nor products of these terms are readily interpretable with respect to the physics at hand, and storage of each of these 4D arrays may be memory-expensive. Additionally, we utilize L2 regression for simplicity and efficiency, as opposed to sparse objective functions such as LASSO or SR3, which are typically used in model discovery frameworks (*30, 40*). For this reason, we refer to our method as function regression, instead of model discovery, as it utilizes a set library constructed from prior knowledge of the physics problem at hand. Some of the PDE parameters are unidentifiable given the structure of the underlying PDE and characteristics of the vascular input function. Further, due to the nature of any overdetermined regression problem, the models recovered may not be unique. This is especially the case for model discovery methods which utilize L1 or SR3, as the discovered model strongly depends on the sparsity parameters used to enforce parsimony. While the present methods may not yield unique results for each individual spatial window, these results consistently indicate the presence of directed transport within tumors and allow for the accurate measurement of transport parameters in vitro and within living tissue.

## Conclusion

In this work, we present a data processing framework tailored specifically to the unique needs of DCE-MRI for studying fluid dynamics in living tissues. These developments were fueled by our interest in understanding interstitial fluid flow and transport, as a key contributing factor influencing cancer biology, progression and response to therapy. Given the lack of tools in this area, our contribution will be of significant interest to the community. In our strategy, we employ approaches from data science and integrate them into DCE-MRI data analysis framework, which allowed us to process and analyze noisy data rapidly *in situ*. Importantly, in an *in vivo* mouse study, we non-invasively capture individual variation in tumor perfusion and interstitial flow over time, and find that perfusion estimated by this non-invasive method are consistent with measures of perfusion and vascular density measured from tissue histology. Finally, we find that fluid velocity magnitude in brain and breast cancers differ significantly, finding the measured velocity to be greater in breast than in brain. Held together, these results strongly support use of our method in measuring both variation across individuals, and between diseases of different origins, providing a novel method for studying the underlying physiology, and demonstrating the application of this novel methodology to routinely collected clinical imaging.

## Acknowledgments

The authors would like to thank the clinicians and researchers who contributed to the creation of the Quantitative Imaging Network Breast-02 and Prostate datasets, and the City of Hope neuro-oncology program. We especially thank all the patients who voluntarily participated in the QIN studies, and their families, for their exemplary strength and generosity. We could not have performed this research without you.

## IRB Statement

The study of publicly available data and retrospective analysis of City of Hope patient data was approved with a waiver of consent by the local Institutional Review Board Protocol 15286.

## Author contributions

Conceptualization: RTW, JRC, JMM, RCR

Methodology: RTW, MW, JCR, RCR, JMM, ABB

Investigation: RTW, CCE, MW

Visualization: RTW, CCE, JRC, JMM, RCR

Funding acquisition: RCR, CCB, JMM

Project administration: RCR, JMM, CCB

Supervision: RCR, JMM, RTW, CCB

Writing – original draft: RTW

Writing – review & editing: RTW, CCE, JCR, MW, MG, CCB, ABB, CS, JMM, RCR

## Data and materials availability

The raw data for analysis, forward models for in silico validation, and raw-output of the LCFR-processed data shown in this work are provided for review. All data and code will be made readily available to readers upon request to the corresponding authors.

## Supplementary Materials

### Materials and Methods

#### In silico forward methods

Forward PDE models of advection, diffusion, and source of contrast in tissue are implemented using the 2-dimensional finite difference method on a 64 mm x 64 mm grid, with Δ*x* = Δ*y* = 1mm, with explicit time stepping. The boundary conditions are Dirichlet such that 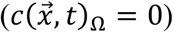

**Table S1:** Description of in silico simulation parameters.

| Simulation | Diffusion Coefficient | Velocity Field | Source Dynamics | Explicit FDM dt |
| --- | --- | --- | --- | --- |
| Divergent Flow | 0.75 mm<sup>2</sup>/s | $u_x = \text{sqrt}(2) \text{ sign}(x) / 2 \text{ mm/s}$<br>$u_y = \text{sqrt}(2) \text{ sign}(y) / 2 \text{ mm/s}$ | None | 1/8 s |
| Poiseuille Shear | 0.75 mm<sup>2</sup>/s | $u_y = 0$<br>$u_x = 2.5 (1-y)^2 / 32^2$ | None | 1/8 s |
| Extended Tofts-Kety | 0 | 0 | K<sub>trans</sub> (1/s):<br>Q1 = 0.1<br>Q2 = 0.2<br>Q3 = 0.4<br>Q4 = 1<br>v<sub>e</sub>: 0.5<br>v<sub>p</sub> = 0.05 | 1/16 min |

#### Collagen-HA in vitro validation

Collagen hydrogels composed of 0.2% collagen (Corning) were crosslinked using 0.4% photo-crosslinkable HA for 45 seconds. Each in vitro phantom consists of 350uL of gel pipetted into a 12mm tissue culture insert (Millipore, Burlington, MA). Prior to imaging, 100uL of 1X PBS was applied below the gel, in a collection chamber. To induce flow through the gel, 300uL of a 1:100 dilution of Gd-DTPA (BioPal, Worcester, MA) in 1X PBS was administered atop the gel. The contrast front velocity was measured by calculating the number of pixels contrast traveled through the gel during the duration of MR imaging (Fig 2, A). Dynamic contrast enhancement imaging parameters match in vivo parameters (See Methods: Magnetic Resonance Imaging, 3D DCE). A single T10 map was acquired through VFA methods (See Methods: Magnetic Resonance Imaging, 3D DCE), across 3 replicates, and the mean T10 value within the gel was used for the T10 value across the 3 replicates to calculate the concentration of Gd-DTPA within the hydrogel.

**Table S2:**
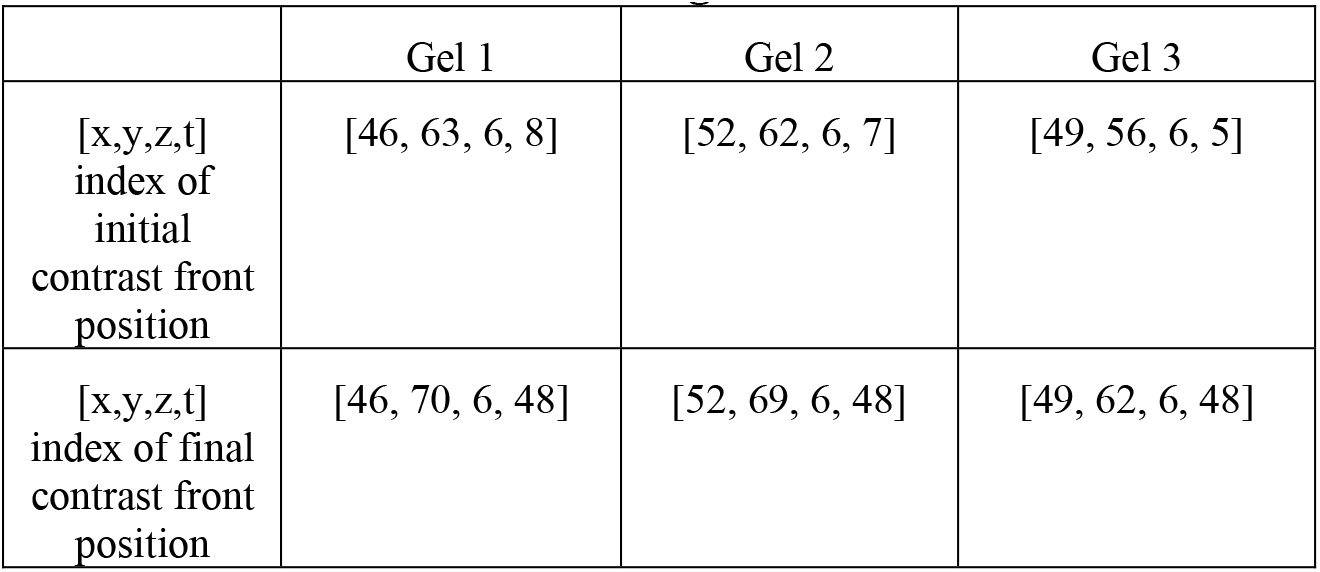
Location of contrast agent front selection.

#### Description of in vivo animal studies

All animal procedures were reviewed and approved by the Institutional Animal Care and Use Committee in accordance with the NIH guidelines for animal research (Institutional Animal Care and Use Committee protocol number 20-146). Three month old, male C57Bl/6 mice (n=6, ∼25g) were purchased from Charles River. The animals were housed in a room maintained at 20-23 °C, 45-55% relative humidity, and a 14-h/12-h light/dark cycle with access to standard laboratory chow and water until the experiment. Mice were injected with 100,000 green fluorescent protein (GFP)-expressing Gl261 (NCI-DTP Cat# Glioma 261, RRID:CVCL_Y003) cells in the cortex and allowed to grow for 14 days. No control group was used in this study as our primary outcome was to correlate flow and histological outcomes within a single animal. Following inoculation, mice were provided wet food and hydrogel for recovery, injected with 4mg/kg ketoprofen for 48 hours post inoculation, and weighed every other day for the duration of the study. Mice were allowed to live in group housing, with a max of four mice per cage. Mice were separated if wounds appeared indicating aggressive behavior.

#### Description of cell lines used

Lentivirus conferring expression of green fluorescent protein (GFP) under puromycin antibiotic selection was a generous gift from the laboratory of Dr. Kevin Janes. Murine GL261, which were originally derived using C57BL/6 male mice (NCI-DTP Cat# Glioma 261, RRID:CVCL_Y003) (*41*), were serially transduced with GFP lentivirus and purified by selection with 2 μg/mL puromycin (Thermo Fisher A1113803). Cells were maintained at 37 °C and 5.2% CO2 for at least three passages after thaw with DMEM + 10% Fetal bovine serum (ThermoFisher, Gibco). Cells were resuspended at a concentration of 20,000 cells/uL in serum free media for tumor implantation. Our cell lines have not undergone cell authentication.

#### Preclinical magnetic resonance imaging

Following anesthesia, a catheter was inserted into the lateral tail vein. Mice were imaged with a 9.4T small animal MRI (Bruker, Ettingen, Germany) equipped with a 20mm RF surface coil. Two consecutive T2-weighted images were collected to verify tumor presence. The first image has high SNR to aid with MRI:IHC alignment for cryosectioning; the second T2-weighted image has isometric voxels for registration to isotropic T1 images. T1 mapping was performed to collect baseline intensity, followed by a 3D DCE T1-weighted FLASH sequence. Six pre-contrast images were acquired before injecting gadolinium (0.2mL/kg, BioPal). A T1-weighted post-contrast image was acquired to confirm contrast enhancement. An extensive list of imaging parameters are included in **Table S1**. After completion of MRI, mice were injected with Evans Blue at a concentration of (1.6mL/kg) which was allowed to circulate overnight before tissue harvesting.

**Table S3:** Preclinical and hydrogel MRI imaging parameters.

|  | 1. T2-weighted | 2. T2-Weighted | 3. T1 mapping | 4. T1-weighted DCE | 5. T1-weighted |
| --- | --- | --- | --- | --- | --- |
| Sequence | RARE | RARE | RARE with varied TR | FLASH | FLASH |
| TE/TR | 40ms/2600ms | 40ms/2600ms | TE: 7ms<br>TR: 5500, 3000, 1500, 800, 500, 300 ms | 4ms/21ms | 3ms/180ms |
| Number of Slices | 16 | 12 | 12 | 1 (3D Imaging) | 16 |
| Slice Thickness | 400 um | 200 um | 200 um | 200 um | 400 um |
| Rare Factor | 8 | 8 | 2 | n/a | n/a |
| Flip angle | 90,180 | 90/180 | 90/180 | 25 | 70 |
| FOV | 19.2x19.2 mm | 19.2 x 19.2 mm | 19.2 x 19.2 mm | 19.2 x 19.2 x 12 mm | 19.2 x 19.2 mm |
| Matrix Size | 192 x 192 | 96 x 96 | 96 x 96 | 96 x 96 x 12 | 192 x 192 |
| Repetitions | 1 | 1 | 1 | 48 | 1 |
| Averages | 9 | 18 | 18 | 1 | 7 |
| Time of Acquisition | 9 min 21 sec | 9 min 21 sec | 9 min 16 sec | 24 min 11 sec | 3 min 1 sec |

#### Tissue harvest and immunohistochemistry

Mice were euthanized and transcardially perfused with 4% PFA and ice cold 1x PBS. Brains were post-fixed in PFA for 18 hours, and placed in 30% sucrose until complete submersion. Afterwards, brains were placed in molds of OCT at -80°C and sectioned at 12um on a cryostat. T2-weighted MRI images were used as a guide during sectioning to inform which MRI slice corresponded to the collected cryosectioned slice. Structural features (i.e ventricles, white matter, tumor shape) were used as visual aides to confirm location, and compared to the corresponding sagittal MRI slice. Brain sections were stained for DAPI (ThermoFisher) and imaged at 20X on an VS200 Olympus Slide Scanner (Olympus).

#### Immunohistochemistry comparison to MRI

To visually compare MRI to histology, all IHC stains were first loaded into MATLAB, and then down-sampled to 0.2 mm in-plane resolution to match the resolution of isotropic MRI. This down-sampling was done using the *blockproc* function in MATLAB, summing the total intensity of sub-pixels within the larger super-pixels so as to maintain the total image intensity. A ROI was drawn around the tumor region of the DAPI stain, using the *drawfreehand* function in MATLAB, as well as on the corresponding slice of post-contrast DCE-MRI.

#### Immunohistochemistry Evans Blue coverage calculation

Evans Blue stains from individual animals were processed using FIJI image processing software to calculate percent tumor coverage. First, a threshold was applied, removing the lower 95^th^ percentile of the image intensity. A mask of the tumor was draw on the DAPI stain to demarcate the tumor using FIJI’s “mask” functionality. This mask was then applied to the Evans Blue stain, and the percent of voxels greater than the 95^th^ percentile intensity within the tumor ROI was reported. The percent tumor Evans Blue coverage was then taken and compared to the mean tumor intensity value of *ξ*_*trans*_ on an individual animal basis, and the correlation and significance of the correlation was determined using a Pearson correlation test.

#### Clinical imaging: QIN BREAST-02

The study of publicly available data and was approved by the local Institutional Review Board Protocol 15286. All 13 patients from the QIN BREAST-02 dataset, provided by The Cancer Imaging Archive (TCIA), were analyzed using LCFR. All patients (female, 18+) were diagnosed with invasive breast cancer, with lesion size > 1cm. Images used were while all patients were treatment-naïve, though all patients received treatment after initial imaging. The multi-flip T1 map and DCE sequences, acquired on Phillips 3T scanners located at both Vanderbilt University Medical Center, and University of Chicago, were used in this study (*33*). Image acquisition details can be found on the TCIA website for the BREAST-02 study.

#### Clinical imaging: Glioblastoma patients

The retrospective study of City of Hope patient data was approved by the local Institutional Review Board Protocol 15286. 20 patients who came to City of Hope National Cancer Center in Duarte, CA, for advanced imaging after resection of glioblastoma between January 2020, and July 2022, with residual enhancing lesion size > 1cm underwent DCE-MRI. Patients underwent imaging between 3 and 135 days post-resection (mean = 32.3, median 28, SD = 30.7). Scans were performed on a 3T Siemens scanner. The DCE scan consisted of 3D FLASH sequence with prior variable-flip angle T1-mapping. Variable Flip angles were acquired at 2,5, and 10 degrees, repetition time = 9.3 ms, and echo time of 4.29 ms. The dynamic scan was performed with 50 phases at 6 second temporal resolution, flip angle = 15 degrees, repetition time = 9.3 ms, and echo time of 4.29 ms. 8 mL of Gadovist (Bayer, Whippany, NJ) was administered during the sequence. The size of the imaging FOV was 192 x 132 in-plane (1.46 mm resolution), and 16 slices with 5 mm slice thickness, identical between the dynamic scan and variable flip angle scans.

**Table S4:** GBM patient age and sex.

| Unique Patient Identifier (UPN) | Patient age at imaging (years) | Patient sex |
| --- | --- | --- |
| 1 | 58 | F |
| 2 | 47 | F |
| 3 | 47 | F |
| 4 | 52 | M |
| 5 | 42 | M |
| 6 | 60 | M |
| 7 | 54 | M |
| 8 | 38 | F |
| 9 | 40 | F |
| 10 | 36 | M |
| 13 | 19 | M |
| 14 | 71 | M |
| 15 | 57 | F |
| 16 | 28 | M |
| 17 | 40 | F |
| 18 | 59 | F |
| 19 | 55 | M |
| 20 | 69 | F |
| 21 | 34 | F |
| 22 | 37 | M |

#### Statistical testing

All statistical analyses and tests were performed using GraphPad Prism 9 (GraphPad Software, San Diego, CA).

#### DCE-MRI pre-processing

Each of the variable flip angle images and dynamic T1-weighted images are rigidly registered to the first dynamic T_1_ image. From images acquired using variable flip angles, α, a T_10_ map for each individual was is calculated by regression (*42*):

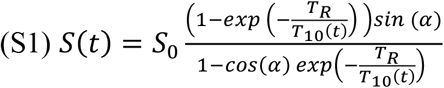

Where S is the measured signal intensity, S_0_ is baseline non-contrast-enhanced signal intensity, TR is the repetition time, T_10_ is the native T_1_ relaxation time of the tissue, and *α* is the flip angle. From the calculated T_10_ map (*43*), and the relaxivity (r_1_) of the contrast agent (Gadovist, 3.7s^-1^mM^-1^ for 3T, and 3.3 7s^-1^mM^-1^for 7T (*44*)), the sequential T_1_-weighted images are converted into a spatiotemporal map of contrast agent concentration (*45*) by

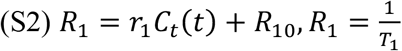

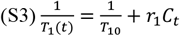

Where R_1_ is the inverse of the T_1_-relaxation time, and R_10_ is the baseline relaxation, or the inverse of the baseline T_10_ relaxation time.

After calculation of contrast agent concentration, the local contrast bolus arrival time (BAT) is calculated by bilinear regression (*46*). Briefly, a sub-set of each individual voxel’s time-enhancement curve, c_sample_, is considered, from the initial time point to the time point with maximal concentration of contrast agent. The points 0, and t_f_ correspond to the initial and final timepoints, and the point p_1_ corresponds to a time points BAT ∈ (0, t_f_). The BAT is determined to be the value that minimizes the summed square error between the bilinear fit and the data

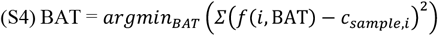

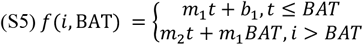

An empirical vascular input function (VIF) is then measured using the automatic-VIF selection algorithm detailed by Singh et al. (*47*). Briefly, this method selects voxels which are rapidly enhancing (BAT *<* 10s), and enhance within the upper 90^th^ percentile of all voxels. The signal intensity is then normalized and scaled to account for partial volume effect and hematocrit. This method is utilized for the CoH GBM patients, the QIN BREAST-02 dataset, and in vivo mouse model. The individual-specific VIF is adjusted for each individual voxel such that the BAT for the VIF matches the estimated BAT of the individual voxel’s enhancement time course, using S4-S5.

#### Weak SINDy for PDEs

Model identification and inversion was performed as follows. Briefly, SINDy identifies dynamics through sparse regression, with the assumption that the time-derivative of the state-space data is a linear combination of polynomial terms of all measured state-space data, and their corresponding spatial and temporal derivatives (*28*):

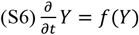

where Y is the measured state-space data. f is a library consisting of linear combinations of analytic functions of Y. In our case, *c*, for contrast agent, is the measured state-variable. In this work, subscript notation is utilized as follows: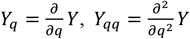. Typically, f consists of the measured state-space data itself, it’s spatial and temporal derivatives, and polynomial combinations of these measurements and their derivatives:

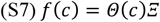

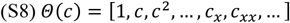

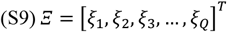

As DCE-MRI is typically modeled as exchange between plasma and extravascular, extra cellular (EES) interstitial space, a BAT-adjusted VIF term and its first derivative are included in the library (*48*). In previous literature, Tofts and Kety parameterized local contrast agent dynamics as

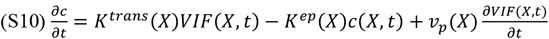

Where, c(X,t) is the contrast agent concentration, K^trans^ is the volumetric flux per unit tissue volume of contrast from vasculature to tissue, K^ep^ is the volumetric return flux per unit tissue volume from tissue to contrast, and v_p_ is the contrast concentration within the plasma compartment, which does not extravasate into the tissue. A practical guide to acquisition of DCE data and its application to this model may be found from Barnes et al. 2012 (*45*). Each of the above terms are included in the function library Θ.

As the VIF may be considered an empirically determined forcing or source function, it is important to include this in the function library. The coefficient in front of the VIF term is mathematically comparable to *K*^*tran*s^, whereas the coefficient in front of the measured concentration is comparable to *K*^*ep*^. Due to potentially high plasma content in high-grade gliomas and other cancers, there may be significant signal contribution from contrast agent within blood plasma. As such, the first temporal derivative of the VIF is also included in the library. The coefficient in front of this term is typically referred to as *v*_*p*_ in a Tofts-Kety framework.

The full function library Θ consists of all terms in the Tofts-Kety model extended by advection and diffusion:

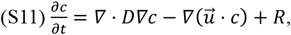

where,

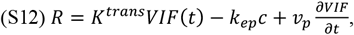

Where D is the diffusion coefficient, and 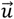 is the contrast agent velocity. The PDE may be expanded to reveal terms as follows:

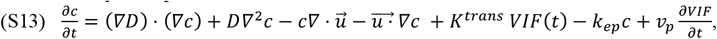

and factored by partial derivatives of terms c(X,t) and VIF(X,t):

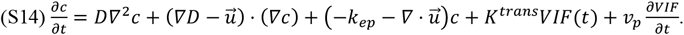

This form of the system may be put in the SINDy framework (S6-S9). Thus, the linear model regression problem takes the form:

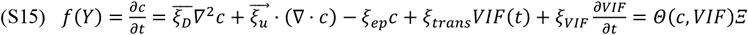

It is assumed that the diffusion is isotropic such that:

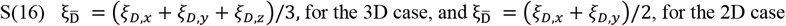

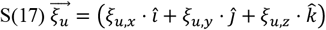

Where 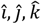 are the standard unit vectors in the (*x, y, z*) directions, respectively.

We do not attempt to separate the contribution to regression coefficients from multiple contributing PDE terms, as they are analytically unidentifiable without estimating the spatial gradient of the diffusivity field D, without using further weak derivatives. As such, we present our results in terms of the regression coefficients *ξ* from the function library, agnostic of the structure of the underlying PDE.

The 3D or 4D data matrix of local, time-resolved concentration of contrast agent, c(x,y,z,t), is acquired either through simulation, or measurement via MRI. MRI data is often noisy in both space and time and may have varying resolutions. To efficiently and accurately perform the regression, rather than utilizing a finite-difference approach for estimating the spatial and temporal derivatives of the measured data, a weak derivative approach is taken to smooth the data. Additional details of the weak PDE regression methodology may be found in Messenger & Bortz (*30*). First, the inner product of (S7) and a given basis function Ψ is taken over the entire spatiotemporal domain on both sides of the equation:

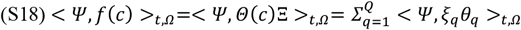

We take the basis functions Ψ to be a tensor product of 1-dimensional polynomial approximations of a 4D Gaussian curve, of the family:

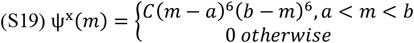

The span of the basis functions [a,b] (in real space/time units (off-grid)) in each dimension is selected using the critical wavelength analysis provided by Messenger and Bortz, 2021. In each dimension, *ψ* is normalized by selection of coefficient *C*, such that

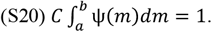

This family of basis functions was selected as it serves to smooth the data as a polynomial approximation to a Gaussian, has analytical expressions for derivatives, and is equal to 0 at the boundary for each of the derivatives required for the construction of the library (for our use case, derivatives up to order 4). Integration by parts is performed on each term in the library, and on the left hand side of the partial differential equation, until the remaining analytic weak derivative form of each term requires no derivatives on the raw data. This methodology effectively transfers the derivatives from the noisy data onto smooth basis functions, while preserving the PDE coefficient in the weights vector Ξ. This is demonstrated in one-dimension as:

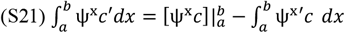

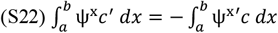

Note that the boundary term vanishes on the boundaries. Practically, Eq S22. is performed using convolution of data *c*, and basis function Ψ or its derivatives. It is assumed that *w* is spatially invariant on a 3x3x3 sample grid of MR data, and over all time, thus, allowing for the weight *w* to be factored from the matrix equation. The vector of weights, Ξ_*n*_, is solved for in parallel using L2 regression on each of the *N* unique 3x3x3 windows of the data matrix, for all time. In the original implementation of SINDy, L1 or SR3 regression is used to enforce sparsity (*28*). This implementation often involves a parameter sweep and selection of an appropriate sparsity-enforcing parameter through Pareto Front analysis (*40*). However, due to the large number of regressions which need to be performed for spatially localized parameterization, the authors opt for L2 regression, avoiding the problem of spatially localized hyperparameter tuning. For each window *n*, the weights *ξ*_*q,n*_ of each weak derivative are calculated:

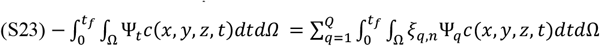

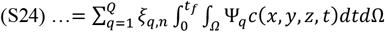

On each individual sliding window *n*, we have

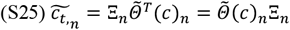

where 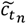 is the weak time-derivative of the measured signal enhancement curve within the window, Ξ_*n*_ is a vector containing the weights *ξ*_*q,n*_, and 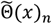is the library containing all corresponding weak spatial and temporal derivatives of the data for sliding window *n* of *N*, where *N* is the total number of sliding windows. This system is solved using L_2_ norm, so as to avoid solving for a localized sparsity enforcing parameter required using the L1, LASSO, or SR3 methodologies:

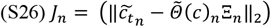

Pseudo-code for the entire localized convolutional function regression algorithm may be found below:

Pseudo-code

1. Initialize *c*(*x, y, z, t*)
2. Select region of interest for VIF, or apply automated VIF selection
3. For each voxel:
  a. Adjust the VIF BAT to match the voxel BAT
4. Assemble library of polynomial terms present in transport equation S15
5. Perform critical wavenumber analysis for each dimension to select the span and shape of the basis functions ψ^*x*^, ψ^*y*^, ψ^*z*^, ψ^*t*^
6. For each dimension, calculate *ψ* and it’s partial derivatives
7. Calculate Ψ (*x, y, z, t*) = *ψ*^*x*^ ⊗ *ψ*^*y*^ ⊗ *ψ*^*z*^ ⊗ *ψ*^*t*^, and all required tensor product combinations for each weak derivative.
8. For each library member, calculate the weak derivative of the data:
  a. θ_*q*_ = -< *c*(*x, y, z, t*), Ψ_*q*_ > _*t*, Ω_
9. Calculate the left-hand side 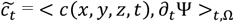= *< c*(*x, y, z, t*), ∂_*t*_Ψ >_*t*, Ω_
10. Subdivide the image into *N* overlapping 3x3x3 windows
11. For each window, *n* of *N*:
  a. Solve the local convolutional function regression using L2 regression for window *n*.
12. For each library term, q of Q:
  a. Overlay heat map of *ξ*_*q*_ on post-contrast MR image

Note, that the above pseudo-code is specific to 3 spatial dimensions. This 3-D implementation was utilized for MRI data consisting of isotropic spatial resolution. An equivalent algorithm was utilized in 2D for data where the z-slice thickness was greater than the in-plane voxel resolution and applied on each individual slice instead of to the entire volume.

The algorithm for this implementation of LCFR is performed in MATLAB 2022b, including the Image and Signal Processing Toolboxes (Natick, MA).

## Supplementary Results

**Supplemental figure 1.**
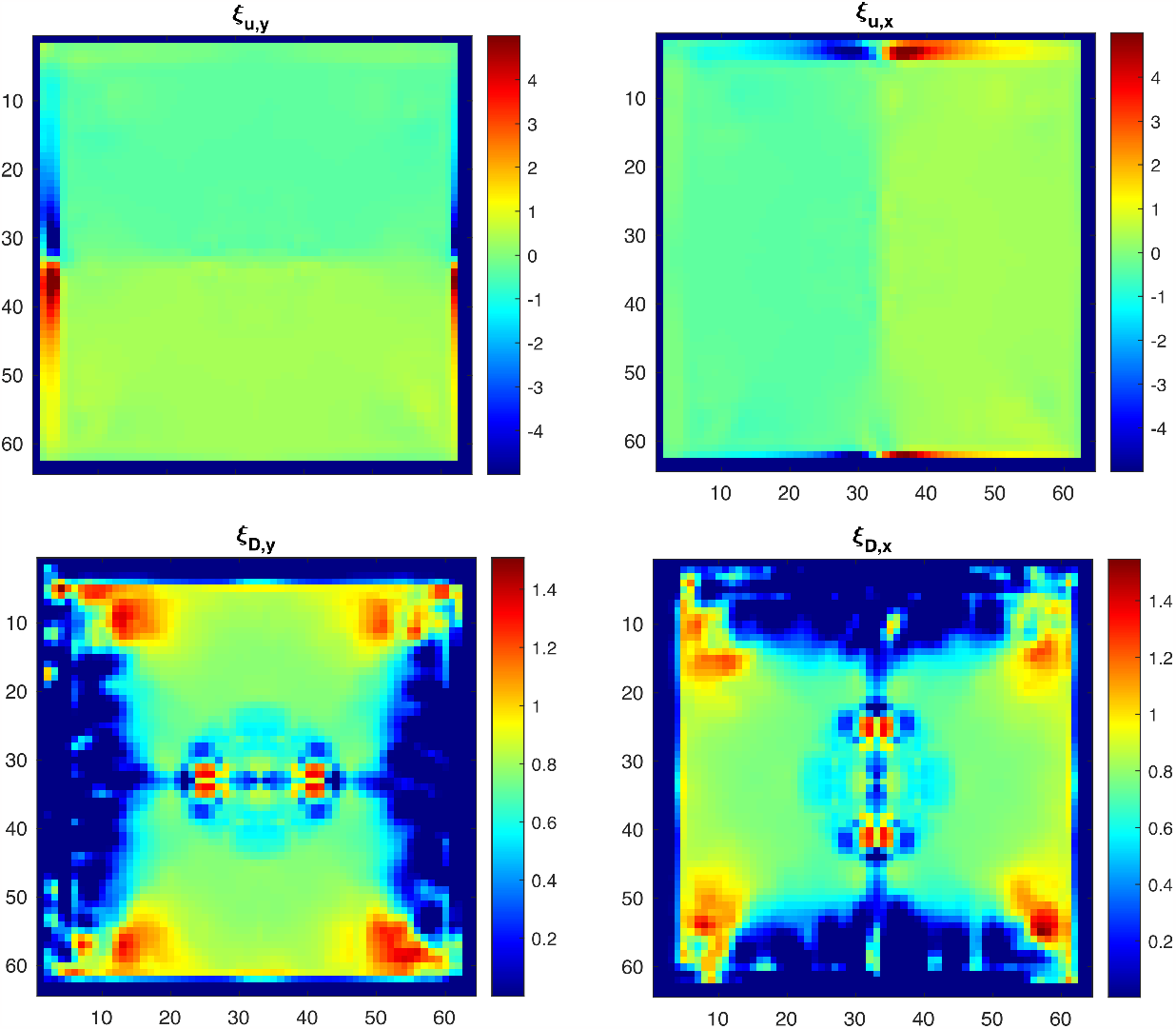
Raw output of outward flow validation. Boundary effects are due to the Gibbs phenomenon resulting from the use of the Fourier transform to perform the convolution of the concentration with the basis functions. Advection accuracy is accurate in regions of strong signal. 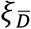 accuracy is also accurate but is more susceptible to edge effects than 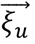.

**Supplemental figure 2.**
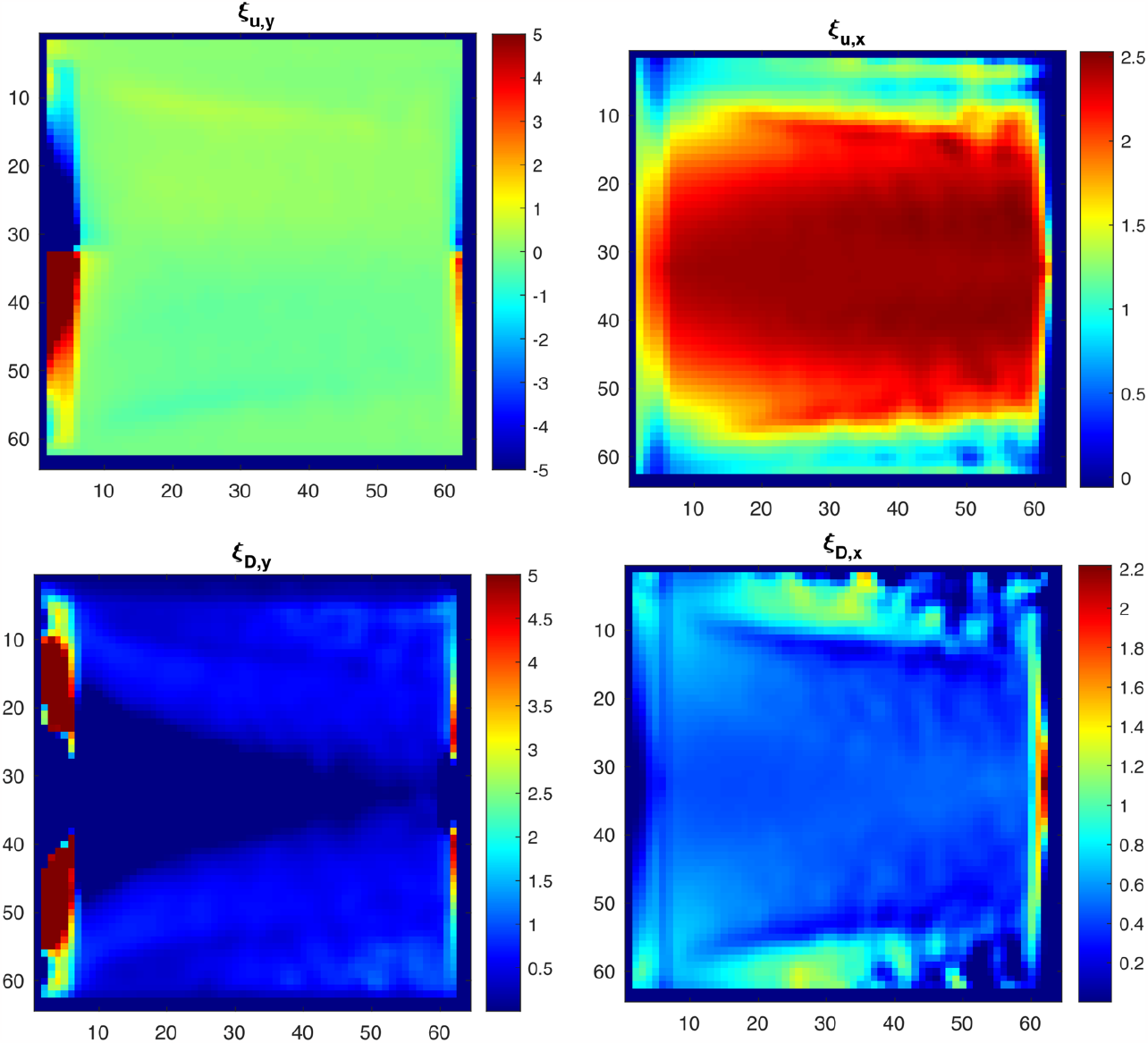
Raw output of shear flow validation. Similar to the outward flow simulation, edge effects are due to the convolution of the concentration with the basis functions and are the result of the Gibbs phenomenon. Edge effects are strongest at initial conditions and boundaries. Accuracy is highest in regions furthest from boundaries, and where contrast was present in high quantities.

**Supplemental figure 3.**
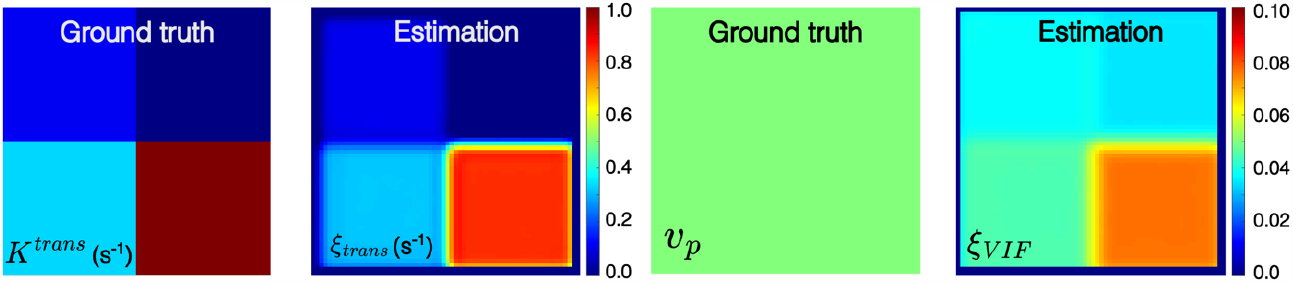
Validation on Tofts-Kety perfusion phantom, where *K*^*trans*^ is varied by quadrant, and *v*_*p*_ is held constant. See Supplemental Table 1 for simulation parameters. LCFR accurately estimates the contrast perfusion rate, *K*^*trans*^, and *v*_*p*_ is accurately estimated, though biased by Ktrans signal. This is largely because *v*_*p*_ and *K*^*trans*^ are often practically non-identifiable.

### Technical Considerations

**Supplemental figure 4.**
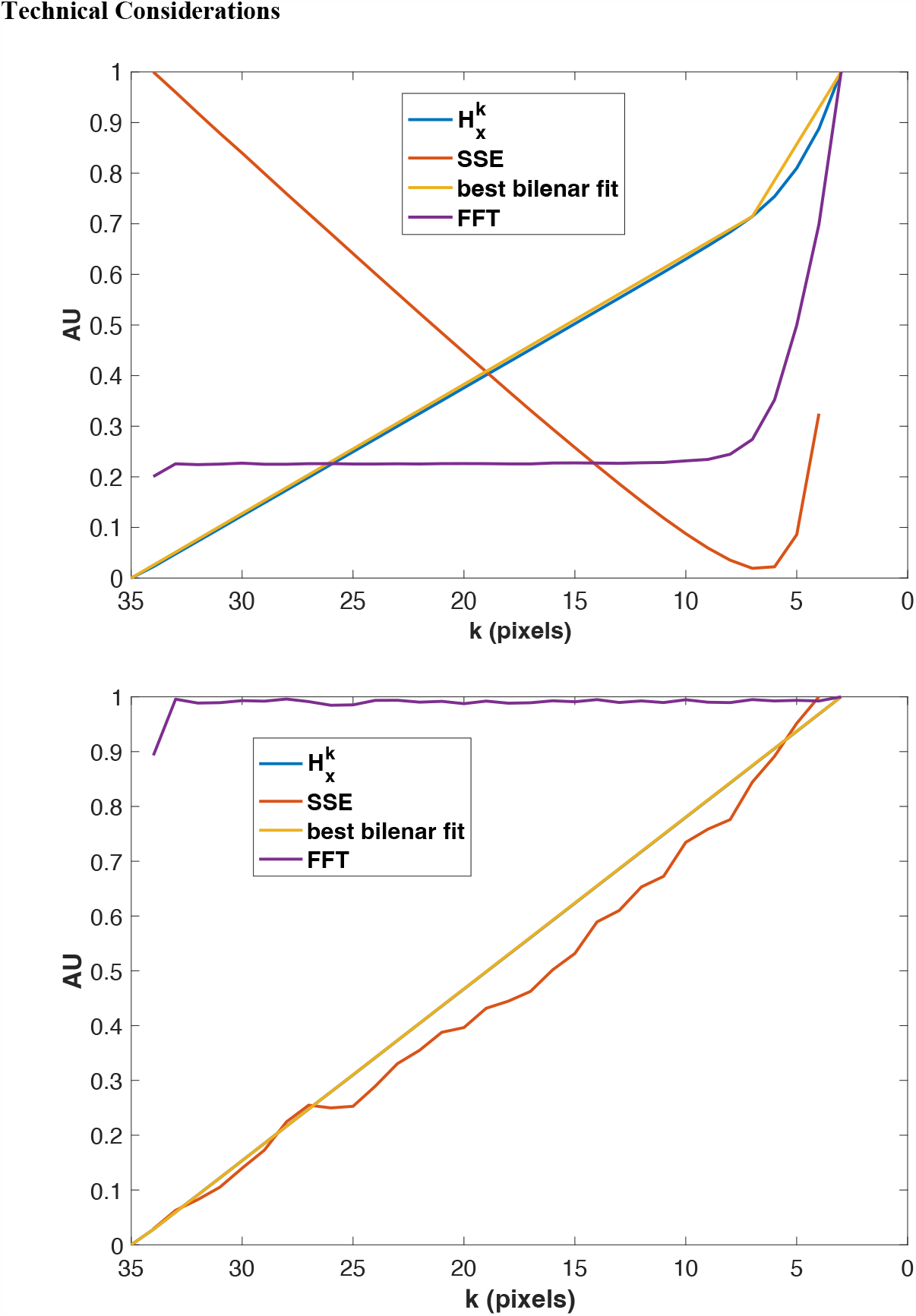
Basis function selection criteria. a) Selection of k* (x-direction) from in silico model of Laminar *Poiseuille* flow, with 1% Gaussian noise added. Note the clear global minimum of the SSE of the bilinear fit selected for k*. b) Identical analysis performed on data with 50% noise added. Note the lack of distinct global minimum in SSE of the bilinear fit. The variance of the Gaussian noise is calculated as σ^2^ = *f* max(*Gd*(*X, t*)), where *f* ∈ (0, 1], or *f* ∈ (0%, 100%] in percent of maximum signal. In Figure 1 (main text), 0.1% noise is added to stabilize the methodology, as the automatic basis function selection algorithm is designed with the expectation of system noise to filter.

#### Basis-function selection

Weak SINDy requires selection of basis functions to smoothly represent discrete data. To automatically select appropriate basis functions, a critical wavenumber, k*, is chosen for each spatial and temporal dimension. This selection is performed with the Fast Fourier Transform of the data in each dimension. Then, the cumulative sum of the FFT, 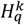 (where *q* is a dimension *x,y,z*, or *t*) is calculated, and approximated by bilinear-fit *b*(*k*),

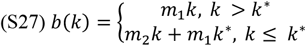

where k* is the critical wavenumber which minimizes the SSE between *b*(*k*) and 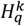. A basis function, in the form of a polynomial low-pass filter (Supplemental Equation 19), is selected such that frequencies greater than *k*^∗^ are filtered out by a factor of 10^−9^ upon convolution of the function with the data. The factor of 10^−9^ was selected empirically, as it provides a balance between noise filtration and minimizing the span of the basis function.

This methodology is useful for MRI data, as it enables the filtering of noise introduced during image acquisition. However, the FFT approach carries with it two primary limitations. First, high frequencies are filtered out from analysis because the resolution of the basis-functions are necessarily lower than the resolution of the image, meaning fine-scale features of the dynamics are filtered and removed from analysis. Second, this method cannot work with data which are too sparse to represent with smooth functions. This second limitation means that weak SINDy cannot be faithfully applied to data which is has fewer samples (*n*_*samples,d*_) less than the polynomial order chosen (*n*_*samples,d*_ < *p -* 1, where *p* is the polynomial order), or where the SSE of the bilinear fit of 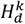 does not have a distinct global minimum. In practice, we have found that this theoretical limitation is insufficient, and that roughly 2*p* datapoints are required. We only present the use of this methodology in cases where there is a distinct separation between signal and noise (through critical wavenumber *k*^∗^ calculation), and where there are sufficient dimensional samples to be accurately represented by a 6^th^ order polynomial. When the z-slice thickness is more than 2x the in-plane resolution, the 2D method is performed on each individual slice. This 2D methodology may also be applied when the number of z-slices is less than 12 (when using 6^th^ order polynomials).

#### Convolutional implementation considerations

Further limitations arise from edge effects due to Gibbs phenomenon at edges, after convolution of the data with the basis functions. Convolution is performed practically using multiplication in k-space after performing an FFT on the data and basis function. These effects are first mitigated by padding with the edge-values using the *padarray* function in MATLAB (Natick, MA), with the ‘replicate’ option set to ‘both’. Second, edge values are removed from the results where Gibbs phenomenon are most prominent.

#### Model assumptions and applicability

This methodology is built on the assumption that contrast agent is present in the tumor interstitium in sufficient quantities to be detected by MR. This assumption only holds for regions where there is sufficient blood supply or leaky vasculature to supply the tissue with contrast agent. As such, the resulting function regression does not correlate strongly with the data in regions with low concentration of contrast agent leakage. Masks were drawn to only including tumor tissue, and to avoiding healthy tissue and vasculature, so as to not bias the numerical analysis towards regions where the model assumptions do not apply.

**Supplemental figure 5.**
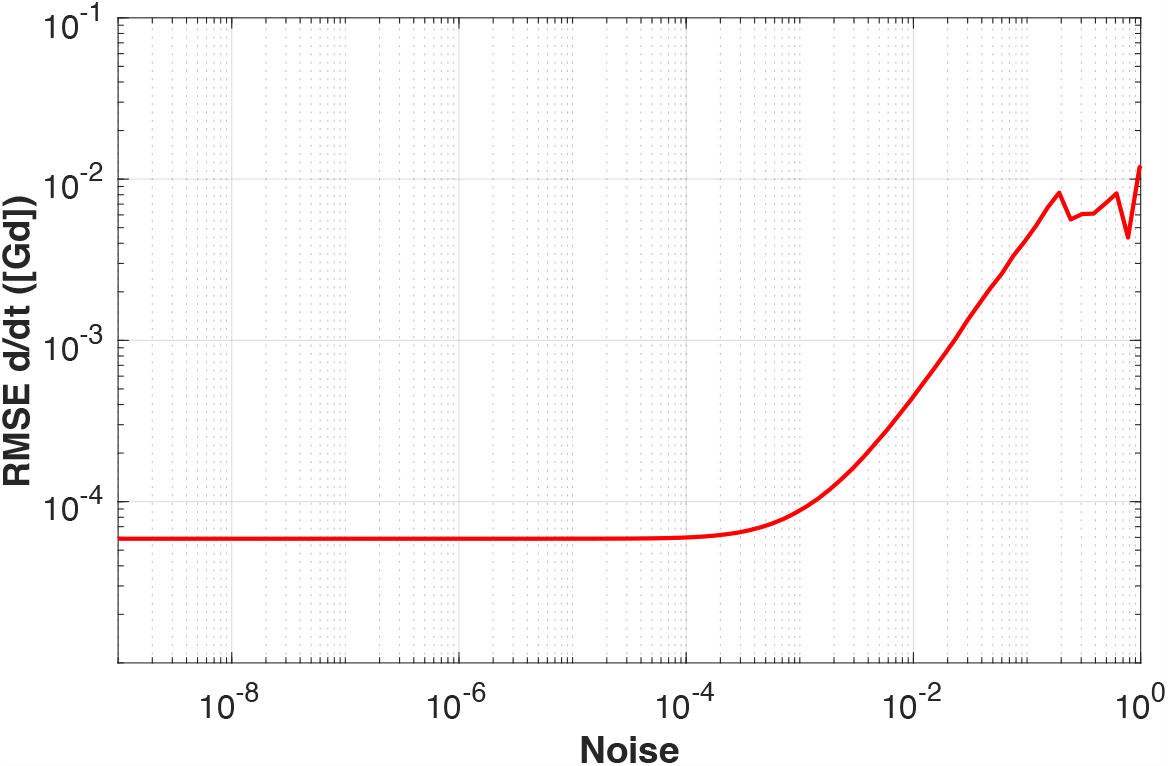
Regression error vs. added Gaussian noise for in silico *Poiseuille* flow. Root mean square error (RMSE) is the regression reconstruction error with respect to the observed weak time-derivative, and the variance of the noise is calculated as σ^2^ = *f* max ([*Gd*(*X, t*)]), where *f* ∈ (0,1]. The RMSE converges to a minimum at approximately *f <* 1*E -* 3, and the reconstruction loses coherence with the original observations at approximately *f* > 1*E -* 1. Our method of weak function regression is robust to noise up to roughly 10% of the maximum signal measured.

#### Limitations due to T_10_-mapping

Model discovery techniques rely on the assumption that the state variable measured is the same variable which is being modeled. The objective of this methodology is to estimate the transport parameters of interstitial fluid, thus the present methodology relies on accurate quantitation of spatiotemporally resolved Gadolinium concentration. As such, we cannot rely on raw MR signal intensity measurements alone. In T_1_-weighted MRI, the T_1_ relaxation time is a non-linear function of contrast agent concentration, *c*:

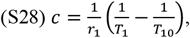

where *r*_*1*_ is the contrast agent relaxivity, *T*_*1*_ is the *T*_*1*_ relaxation time upon interaction with the contrast agent, and *T*_*1*0_ is the native tissue *T*_1_ relaxation time without the presence of contrast agent. As such, a *T*_10_ map must be acquired prior to contrast injection, typically through the variable flip angle (VFA) method or variable TR (VTR) method (*49*). This allows for the measured T_1_-weighted signal intensity to be corrected for T_10_ inhomogeneity of the tissue, and the true concentration to be estimated in space and time. In this work, we present imaging results only where a T_10_ map was acquired correctly. In some cases, it has been shown that a singular literature T_10_ value may be used to estimate contrast agent concentration (*50*). While this practice may be admissible in practice for ODEs parameter estimation of contrast agent transport, it results in spurious spatial concentration gradients required for PDEs parameter estimation. This effect can be seen in the QIN Prostate dataset, where a literature T_10_ value of 1600 ms was used. **Figure S5** depicts the predicted direction of the vector 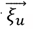 clearly demonstrating a strong dependence on strong gradients in the image.

**Supplemental figure 6:**
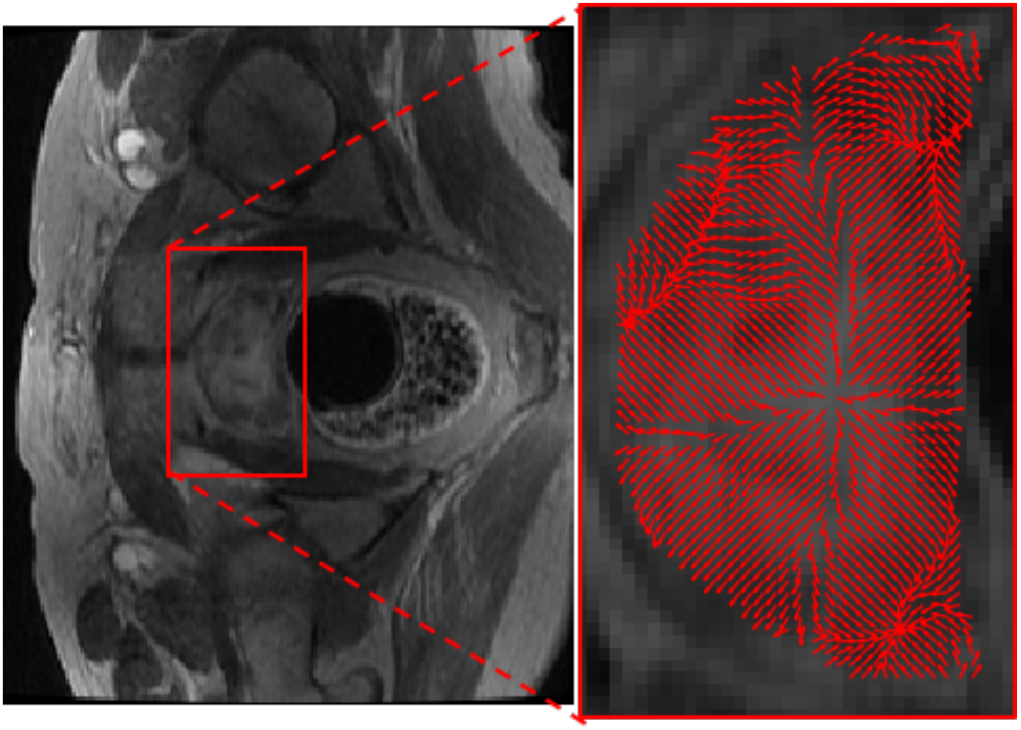
Demonstration of technique on dataset without T10-map. The direction of 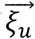 strongly correlates with anatomical gradients along the capsule of the prostate, due to inability to calculate the T10 map directly from a properly-acquired VFA MRI sequence. This dataset was acquired with variable flip angles, but the flip angle scan also had varying repetition time, leading to an inability to calculate a proper T1-map, forcing a literature value to be used. As such, this data was not included in the analysis. All other figures shown in this work utilize a VFA- or VTR-acquired *T*_10_ map to calculate the proper concentration, preventing the occurrence of spurious image gradients due to variations in native *T*_10_ from the analysis.

**Supplemental figure 7:**
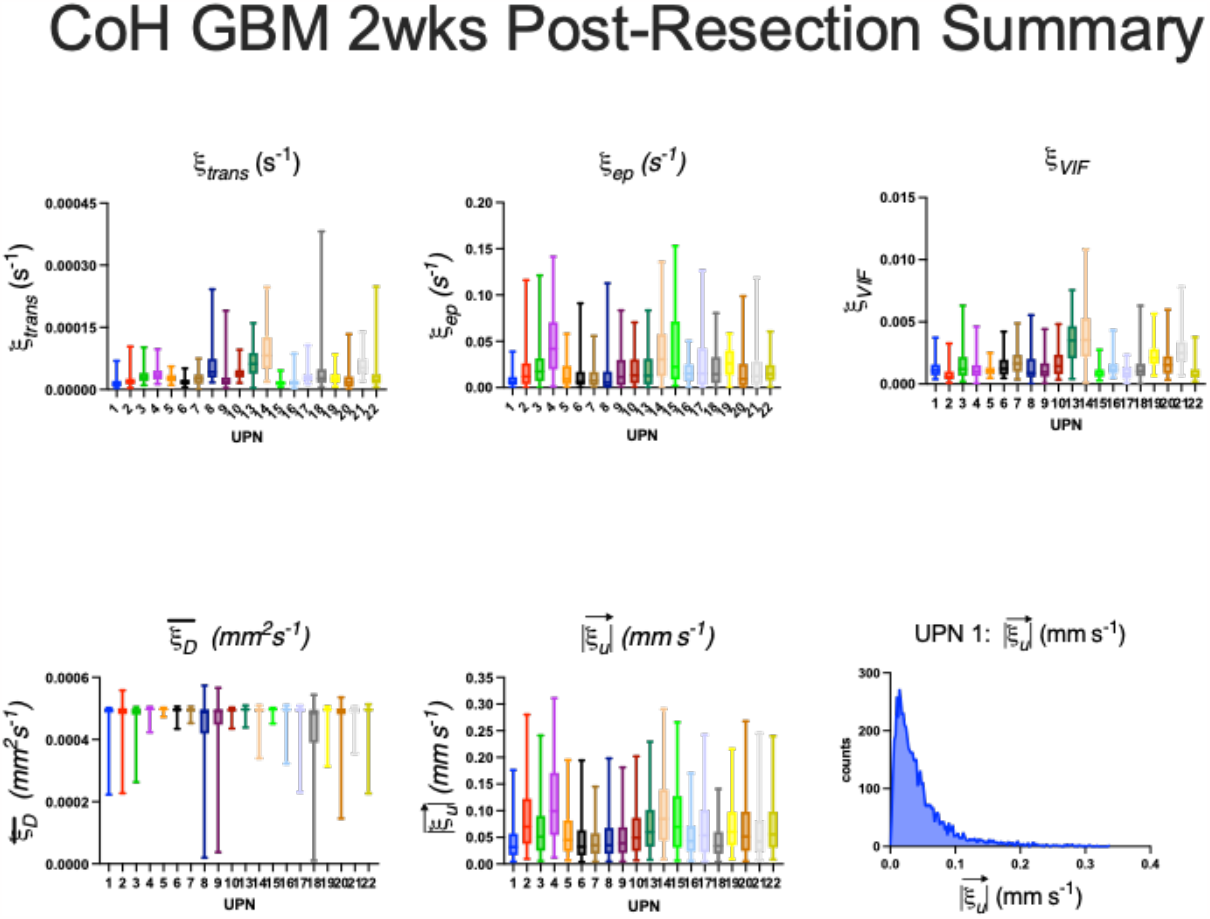
Summary of LCFR coefficients in the CoH GBM Cohort. The in-plane resolution of the voxels was 1.46 mm, and the slice-thickness was 5 mm. As the slice thickness was more than 2 times the in-plane resolution, the method was applied in 2 dimensions, neglecting the z-component of velocity- and diffusion-associated parameters. The results shown here, and in the tables, below are the results of analysis over a hand-drawn mask, and the logical intersection where the R^2^ value of the method with respect to the weak-time derivative of the data is R^2^ > 0.5.

**Table S5:** CoH GBM Cohort Summary, Parameter Means.

| UPN | $\vert \vec{\xi}_u\vert $ (mm/s) | $\bar{\xi}_D$ (mm<sup>2</sup>/s) | $\xi_{VIF}$ (frac.) | $\xi_{ep}$ (s<sup>-1</sup>) | $\xi_{trans}$ (s<sup>-1</sup>) |
| --- | --- | --- | --- | --- | --- |
| 1 | 4.52 E-2 | 4.76 E-4 | 1.32E-3 | 9.27 E-3 | 1.82 E-5 |
| 2 | 8.99 E-2 | 4.78 E-4 | 8.584 E-4 | 2.40 E-2 | 2.46 E-5 |
| 3 | 6.83 E-2 | 4.73 E-4 | 1.74 E-3 | 2.51 E-2 | 3.55e E-5 |
| 4 | 1.19 E-1 | 4.93 E-4 | 1.29 E-3 | 5.15 E-2 | 3.87e E-5 |
| 5 | 6.04 E-2 | 4.95 E-4 | 1.13 E-3 | 1.62 E-2 | 2.85e E-5 |
| 6 | 4.88 E-2 | 4.904 E-4 | 1.50 E-3 | 1.57 E-2 | 2.02e E-5 |
| 7 | 4.41 E-2 | 4.92 E-4 | 1.85 E-3 | 1.25 E-2 | 2.70e E-5 |
| 8 | 5.23 E-2 | 4.29 E-4 | 1.53 E-3 | 1.61 E-2 | 6.47e E-5 |
| 9 | 5.23 E-2 | 4.44 E-4 | 1.31 E-3 | 2.03 E-2 | 3.28 E-5 |
| 10 | 6.30 E-2 | 4.90 E-4 | 1.79 E-3 | 2.05 E-2 | 4.11 E-5 |
| 13 | 7.52 E-2 | 4.92 E-4 | 3.51 E-3 | 2.13 E-2 | 6.70 E-5 |
| 14 | 1.01 E-1 | 4.84 E-4 | 4.00 E-3 | 3.93 E-2 | 9.60 E-5 |
| 15 | 8.80 E-2 | 4.92 E-4 | 9.76 E-4 | 4.35 E-2 | 1.75 E-5 |
| 16 | 5.41 E-2 | 4.84 E-4 | 1.43 E-3 | 1.78 E-2 | 2.03 E-5 |
| 17 | 7.06 E-2 | 4.79 E-4 | 9.53 E-4 | 2.78 E-2 | 3.45 E-5 |
| 18 | 4.51 E-2 | 4.14 E-4 | 1.47 E-3 | 2.20 E-2 | 6.05 E-5 |
| 19 | 7.38 E-2 | 4.81 E-4 | 2.34 E-3 | 2.66 E-2 | 3.04 E-5 |
| 20 | 7.35 E-2 | 4.68 E-4 | 1.94 E-3 | 1.97 E-2 | 2.94 E-5 |
| 21 | 6.32 E-2 | 4.83 E-4 | 2.86 E-3 | 2.17 E-2 | 5.95 E-5 |
| 22 | 7.40 E-2 | 4.79 E-4 | 1.05 E-3 | 1.85 E-2 | 4.41 E-5 |
| Mean | 6.81E-2 | 4.76 E-4 | 1.741 E-3 | 2.35 E-2 | 3.95 E-5 |

**Table S6:** CoH GBM Cohort Summary, Parameter 95% CIs.

| UPN | $\vert \vec{\xi}_u\vert $ (mm/s) | $\bar{\xi}_D$ (mm<sup>2</sup>/s) | $\xi_{VIF}$ (frac.) | $\xi_{ep}$ (s<sup>-1</sup>) | $\xi_{trans}$ (s<sup>-1</sup>) |
| --- | --- | --- | --- | --- | --- |
| 1 | 4.41E-3 | 6.98E-5 | 1.01E-3 | 1.02E-2 | 1.89E-5 |
| 2 | 7.18E-2 | 7.95E-5 | 8.62E-4 | 4.47E-2 | 2.79E-5 |
| 3 | 6.13E-2 | 6.32E-5 | 1.79E-3 | 2.89E-2 | 2.99E-5 |
| 4 | 8.41E-2 | 2.60E-5 | 1.06E-3 | 4.15E-2 | 2.26E-5 |
| 5 | 5.09E-2 | 2.19E-5 | 5.29E-4 | 1.66E-2 | 1.32E-5 |
| 6 | 4.95E-2 | 3.15E-5 | 1.01E-3 | 2.43E-2 | 1.48E-5 |
| 7 | 3.60E-2 | 1.93E-5 | 1.19E-3 | 1.47E-2 | 1.89E-5 |
| 8 | 5.10E-2 | 1.42E-4 | 1.89E-3 | 2.67E-2 | 6.61E-5 |
| 9 | 4.56E-2 | 1.23E-4 | 1.25E-3 | 2.34E-2 | 5.44E-5 |
| 10 | 5.19E-2 | 2.59E-5 | 1.26E-3 | 2.03E-2 | 2.02E-5 |
| 13 | 5.86E-2 | 2.81E-5 | 2.04E-3 | 2.36E-2 | 4.09E-5 |
| 14 | 7.39E-2 | 5.80E-5 | 3.45E-3 | 3.50E-2 | 6.50E-5 |
| 15 | 7.12E-2 | 3.20E-5 | 7.42E-4 | 4.69E-2 | 1.16E-5 |
| 16 | 4.38E-2 | 5.47E-5 | 9.87E-4 | 1.44E-2 | 2.17E-5 |
| 17 | 6.27E-2 | 6.92E-5 | 6.09E-4 | 3.28E-2 | 3.42E-5 |
| 18 | 3.81E-2 | 1.49E-4 | 1.62E-3 | 2.33E-2 | 1.04E-4 |
| 19 | 5.48E-2 | 5.91E-5 | 1.19E-3 | 1.70E-2 | 2.73E-5 |
| 20 | 6.84E-2 | 9.38E-5 | 2.27E-3 | 2.64E-2 | 5.67E-5 |
| 21 | 6.02E-2 | 4.80e-5 | 1.84E-3 | 2.78E-2 | 3.12E-5 |
| 22 | 6.13E-2 | 7.68e-5 | 9.32E-4 | 1.57E-2 | 9.9E-5 |
| SD of Patient Mean | 1.99E-2 | 2.19E-5 | 8.45E-4 | 1.04E-2 | 2.038E-5 |

**Supplemental figure 8:**
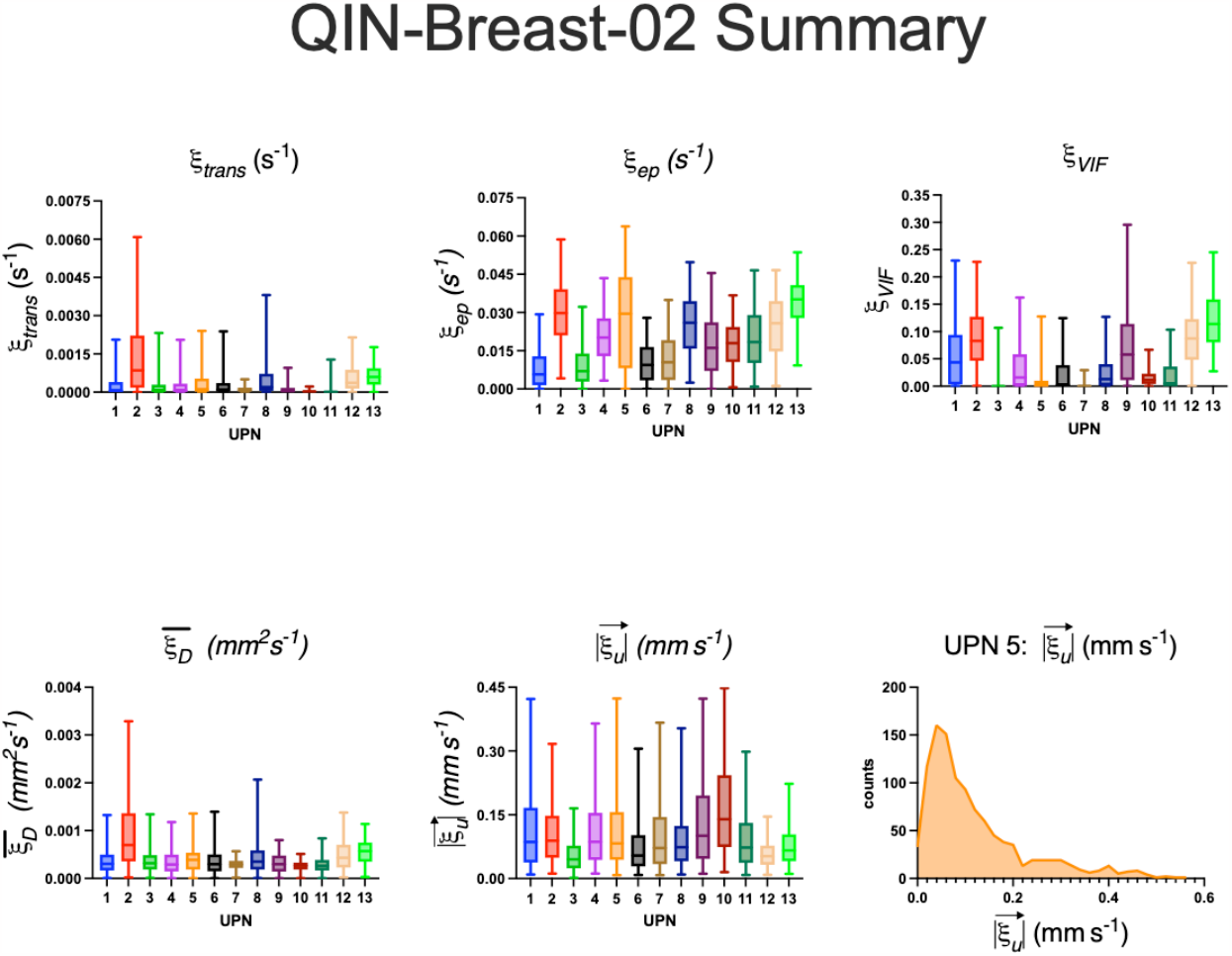
Summary of LCFR coefficients on the QIN Breast-02 DCE MRI dataset. The in-plane resolution of the voxels was 1.33 mm, and the slice-thickness was 5 mm. As the slice thickness was more than 2 times the in-plane resolution, the method was applied in 2 dimensions, neglecting the z-component of velocity- and diffusion-associated parameters. The results shown here, and in the tables below, are the results of analysis within a hand-drawn mask of enhancing tumor, and the logical intersection where the R^2^ value of the method with respect to the weak-time derivative of the data is R^2^ > 0.5.

**Table S7:** QIN Breast-02 Dataset Summary, Parameter Means.

| UPN | $\vert \xi_u\vert $ (mm/s) | $\xi_D$ ($mm^2/s$) | $\xi_{VIF}$ (frac.) | $\xi_{ep}$ ($s^{-1}$) | $\xi_{trans}$ ($s^{-1}$) |
| --- | --- | --- | --- | --- | --- |
| 1 | 1.19E-1 | 4.53 E-4 | 6.06E-2 | 7.99E-3 | 3.39E-4 |
| 2 | 1.09E-1 | 4.83 E-4 | 9.16 E-2 | 3.05E-2 | 1.50 E-3 |
| 3 | 5.67E-2 | 4.49 E-4 | 1.13 E-2 | 9.23E-3 | 3.08 E-4 |
| 4 | 1.12E-1 | 3.97E-4 | 3.48 E-2 | 2.08E-2 | 3.23 E-4 |
| 5 | 1.19E-1 | 4.12E-4 | 1.62 E-2 | 2.79E-2 | 4.27 E-4 |
| 6 | 8.02E-2 | 4.22E-4 | 2.36 E-2 | 1.06E-2 | 3.82 E-4 |
| 7 | 1.04E-1 | 4.45E-4 | 2.93E-3 | 1.20E-2 | 1.41 E-4 |
| 8 | 9.73E-2 | 3.48E-4 | 2.74E-2 | 2.55E-2 | 6.66 E-4 |
| 9 | 1.37E-1 | 4.90E-4 | 7.69E-2 | 1.74E-2 | 1.73 E-4 |
| 10 | 1.70E-1 | 4.81E-4 | 1.65E-2 | 1.76E-2 | 7.35 E-5 |
| 11 | 9.52E-2 | 4.57E-4 | 2.18E-2 | 2.00E-2 | 1.26E-4 |
| 12 | 5.89E-2 | 4.62E-4 | 9.04E-2 | 2.51E-2 | 5.83E-4 |
| 13 | 7.94E-2 | 4.64E-4 | 1.22E-1 | 3.39E-2 | 6.87E-4 |
| Mean | 1.02E-1 | 4.43E-4 | 4.59E-2 | 1.99 E-2 | 4.41E-4 |

**Table S8:** QIN Breast-02 Dataset Summary, Parameter 95% Confidence Interval.

| UPN | $\vert \vec{\xi}_u\vert $ (mm/s) | $\xi_D$ (mm<sup>2</sup>/s) | $\xi_{VIF}$ (frac.) | $\xi_{ep}$ (s<sup>-1</sup>) | $\xi_{trans}$ (s<sup>-1</sup>) |
| --- | --- | --- | --- | --- | --- |
| 1 | 2.68E-3 | 6.10E-6 | 1.69E-3 | 2.14E-4 | 1.32E-5 |
| 2 | 6.66E-4 | 1.93E-6 | 4.94E-4 | 1.13E-4 | 1.41E-5 |
| 3 | 1.32E-3 | 8.03E-6 | 8.68E-4 | 2.61E-4 | 1.70E-5 |
| 4 | 2.06E-3 | 6.02E-6 | 1.03E-3 | 2.39E-4 | 1.30E-5 |
| 5 | 3.23E-3 | 8.88E-6 | 1.07E-3 | 6.13E-4 | 2.00E-5 |
| 6 | 2.42E-3 | 7.24E-6 | 1.18E-3 | 2.74E-4 | 1.92E-5 |
| 7 | 1.69E-3 | 3.99E-6 | 1.37E-4 | 1.84E-4 | 2.53E-6 |
| 8 | 1.86E-3 | 4.88E-6 | 7.92E-4 | 2.86E-4 | 2.37E-5 |
| 9 | 1.35E-3 | 3.05E-6 | 9.44E-4 | 1.49E-4 | 3.63E-6 |
| 10 | 4.04E-3 | 5.65E-6 | 5.80E-4 | 3.29E-4 | 2.60E-6 |
| 11 | 1.47E-3 | 5.31E-6 | 5.90E-4 | 2.38E-4 | 5.96E-6 |
| 12 | 1.62E-3 | 9.93E-6 | 2.58E-3 | 5.61E-4 | 2.49E-5 |
| 13 | 1.73E-3 | 7.20E-6 | 1.73E-3 | 3.31E-4 | 1.48E-5 |
| SD of Patient Mean | 3.10E-2 | 3.97E-5 | 3.81E-2 | 8.41E-3 | 3.76E-4 |

